# Survey of the human proteostasis network: the ubiquitin-proteasome system

**DOI:** 10.64898/2026.03.13.711689

**Authors:** Suzanne Elsasser, Evan T. Powers, Thomas Stoeger, Xiaojing Sui, Roy D. Kurtzbard, Patricia Martínez-Botía, Margaret A. Wangeline, Alejandro Rodriguez Gama, Edward L. Huttlin, Lisa P. Elia, Jeffery W. Kelly, Jason E. Gestwicki, Judith Frydman, Steven Finkbeiner, Eugenia M. Clerico, Richard I. Morimoto, Miguel A. Prado, Alfred C.O. Vertegaal, Kay Hofmann, Daniel Finley

## Abstract

Modification by ubiquitination governs the half-lives of thousands of proteins that are fated for elimination by either the proteasome or autophagy pathways, depending on the intricate architectures of ubiquitin modification. This system mediates quality control for individual proteins, protein complexes, and organelles, as well as myriad purely regulatory functions. Here we provide a comprehensive survey of the ubiquitin-proteasome system (UPS), the scope of which is at present poorly defined. The UPS, with the inclusion of pathways involving ubiquitin-like modifiers, comprises in our estimate over 1430 distinct proteins in humans, a vast set of activities whose collective impact on the biology of the cell is pervasive. The UPS is an integral component of the proteostasis network (PN), the remainder of which we have also surveyed in recent studies. With the addition of molecular chaperones, proteins from autophagy-lysosome pathway, and related activities, the PN includes in total over 3150 components by our estimates. Comprehensive and systematic definition of these pathways should support a range of ongoing investigations in the areas of genomics, proteomics, biochemistry, cell biology, and disease research.

## INTRODUCTION

Modification of target proteins with the small protein ubiquitin, first identified 49 years ago^1,2^, can target proteins for degradation by the proteasome or by autophagy, and also functions in nonproteolytic signaling. These distinct effects on protein fate and function largely reflect the number of bound ubiquitin groups and their arrangement, typically in multiubiquitin chains of varied architecture. The configuration of the substrate-bound ubiquitin chain is the foundational feature of the ubiquitin code, and arises from highly specific utilization of any of eight ubiquitin acceptor sites in ubiquitin itself, seven of these being lysine residues^3^. As originally conceived^4^, the ubiquitin-proteasome system consisted of a small handful of enzymes (**Fig. 1**): ubiquitin-activating enzyme, ubiquitin conjugating enzyme, ubiquitin ligase, a protease later called the proteasome^5^, and a deubiquitinating enzyme. Already by 1985, as ubiquitin-affinity chromatography was refined, it became apparent that some of these biochemical activities actually correspond to families of enzymes^6^, and a stunning multiplicity of ubiquitin ligases was eventually uncovered^7,8^, here estimated to be at least 790 in humans (see Methods). Likewise, ubiquitin recognition was originally attributed to a single receptor on the proteasome^9^, but we currently classify more than 420 proteins as likely “interpreters” of the ubiquitin signal in its many forms, including ubiquitin-like modifiers as discussed below. There is also an unexpected functional diversity of deubiquitinating enzymes, which now number well over 100.

**Figure 1.**
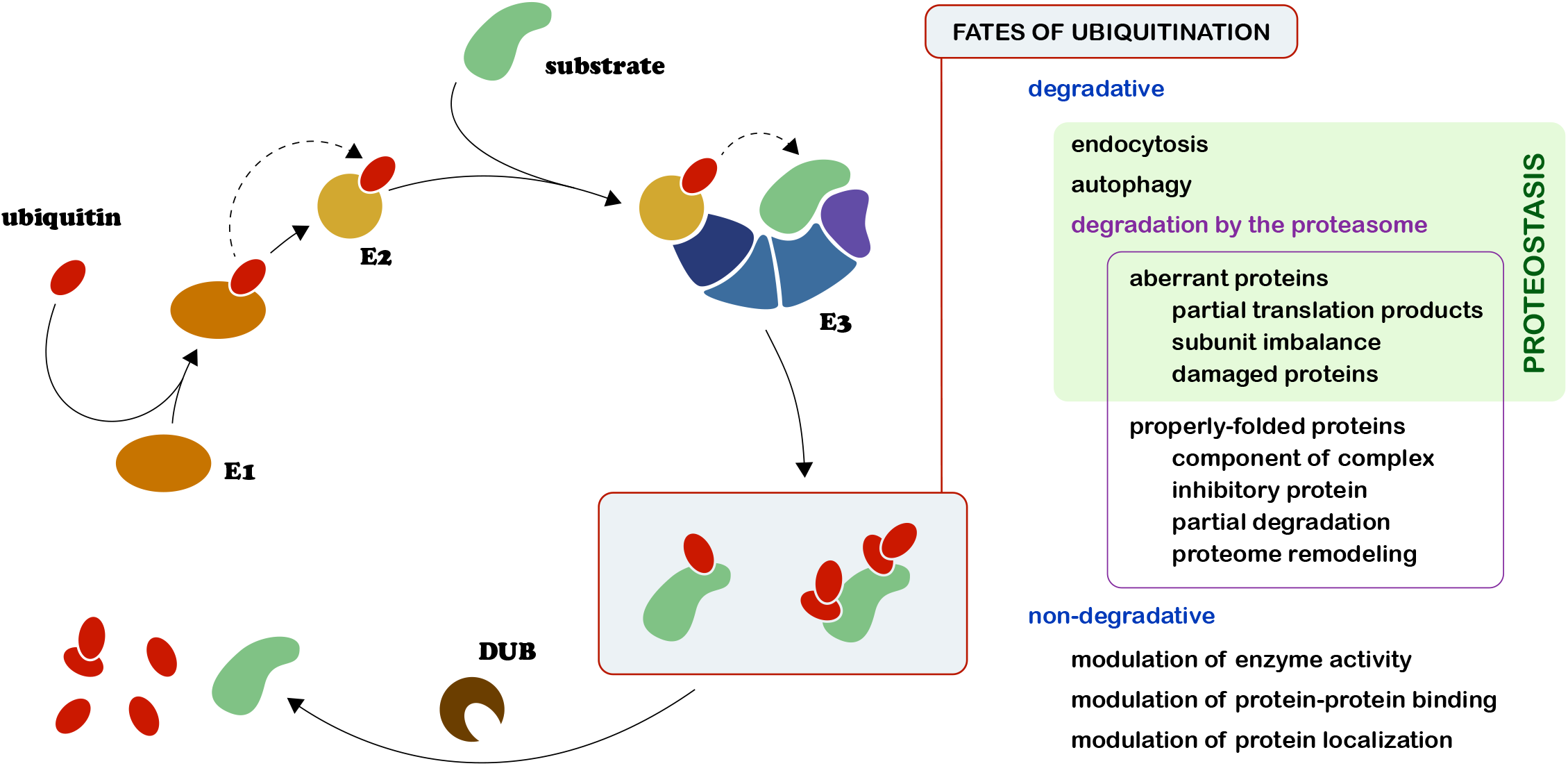
The ubiquitin cycle. ***Left:*** Ubiquitin (red) is activated by E1 (orange) in an ATP-dependent reaction which results in ubiquitin forming a thioester adduct with E1. Ubiquitin is transferred to E2 (gold) in a transthiolation reaction. Charged E2 is recruited by the E3 ligase via an E2 docking element (dark blue). Substrate (green) is recruited by the E3 (blues and purple), and ubiquitin is transferred from E2 to substrate, or from E2 to E3 to substrate, depending on the E3 group. Substrate modification may be reversed by deubiquitinating enzymes (brown). ***Right:*** the fates of substrate ubiquitination, which may include engagement with a number of pathways, resulting in either degradative or non-degradative fates.

With likely UPS members now at 1438 in number, as estimated below, most reviews of the topic have been focused understandably on a specific class of enzymatic activity, often ubiquitin ligases. At the outset of the present study, a systematic accounting of all members of the entire pathway had not been attempted for many years. Therefore, the scope of the UPS is still in this sense lacking in definition, and is not well demarcated as a specific set of gene products. No catalog has been available for querying the entire UPS at the resolution of individual genes, which is a hindrance to many areas of biological research. Given the deep impact of ubiquitination in the biology of the cell, in human health, and in therapeutics^10^, the utility of defining the scope of the UPS in terms of its protein components is evident. For example, genomics and proteomics data now being generated on a tremendous scale are typically filtered through bioinformatic pipelines, or classification systems, based on distinct biochemical systems such as ubiquitination. Proper interpretation of such data is limited by the quality of the assignment of system components, impacting the depth and precision of the biological literature in many ways.

We have recently worked to define the protein components of other biochemical systems that are functionally related to the UPS, namely autophagy, protein synthesis machinery, molecular chaperones, systems for trafficking proteins into and out of organelles, and organelle-specific protein degradation systems (**Fig. 2**)^11,12^. In total, these are parsed as nine main branches that form the proteostasis network, or PN^13^. The PN is defined as the entirety of activities that ensure protein homeostasis, and can be viewed most simply, albeit reductively, as a vast pathway for protein quality control. PN pathways ensure accurate protein synthesis, protein folding, and the removal of damaged, mislocalized, or mutant proteins, and the UPS constitutes a central component of this apparatus. In supporting the elimination of aberrant proteins, it works closely with the molecular chaperone machinery and autophagy. Indeed, many proteins and organelles are targeted to autophagy by ubiquitination^14^.

**Figure 2.**
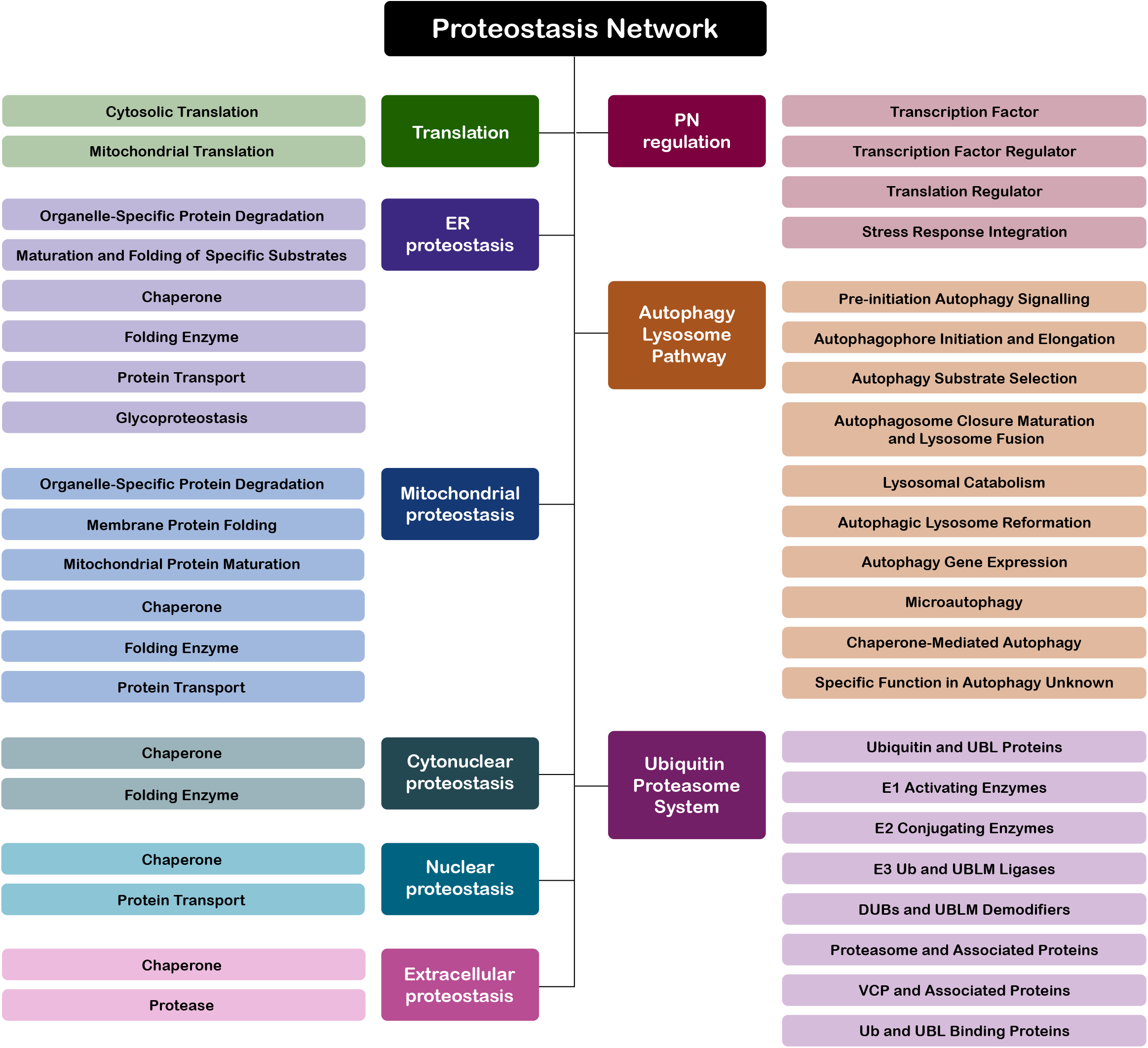
The proteostasis network. The nine Branches of the Proteostasis Network are shown in two center columns. A Branch comprises multiple Classes, shown flanking each Branch. Progressive subdivisions of the Classes include Groups, Types, and Subtypes (not shown; see **Table S1**). Tallies for each of the Branches and Classes are given in **Table S1**.

While the UPS promotes proteostasis by degrading damaged and aberrant proteins, it also degrades properly formed proteins, particularly in response to environmental, developmental, and cell-cycle cues. These more strictly regulatory functions of UPS lie largely outside of proteostasis and the PN per se. However, given the current state of knowledge, we are far from being able to generally classify UPS components as specifically serving either proteostasis or other roles. We therefore aimed to be inclusive of UPS components in our survey, and not to focus specifically on UPS components that support protein or organellar quality control.

For the foreseeable future, any attempt to comprehensively survey the UPS, however carefully done, will carry caveats. For example, hundreds of proteins whose features signify membership in the UPS have not yet been studied in any detail, and the taxonomic weight of their features, such as specific structural domains, is subject to debate in certain instances, as discussed below. Another caveat is that new pathway components presumably remain to be discovered.

To make this survey as accessible as possible, we developed a five-tiered taxonomic system, proceeding from broad to narrow through Branch, Class, Group, Type, and Subtype (**Fig. 3**). The complete survey is presented in tabular form (**Table S1**), and in abbreviated form for easier reading (**Table S2**). The UPS encompasses three Classes that cooperate to write the ubiquitin signal (E1, E2, and E3), one that erases it (deubiquitinating enzymes), one that constitutes the signal itself (ubiquitin and ubiquitin-like proteins), and one that reads it (ubiquitin receptors). The remaining two Classes consist of the molecular machines that determine the fate of proteins carrying the ubiquitin signal (the proteasome and VCP/p97/Cdc48).

**Figure 3.**
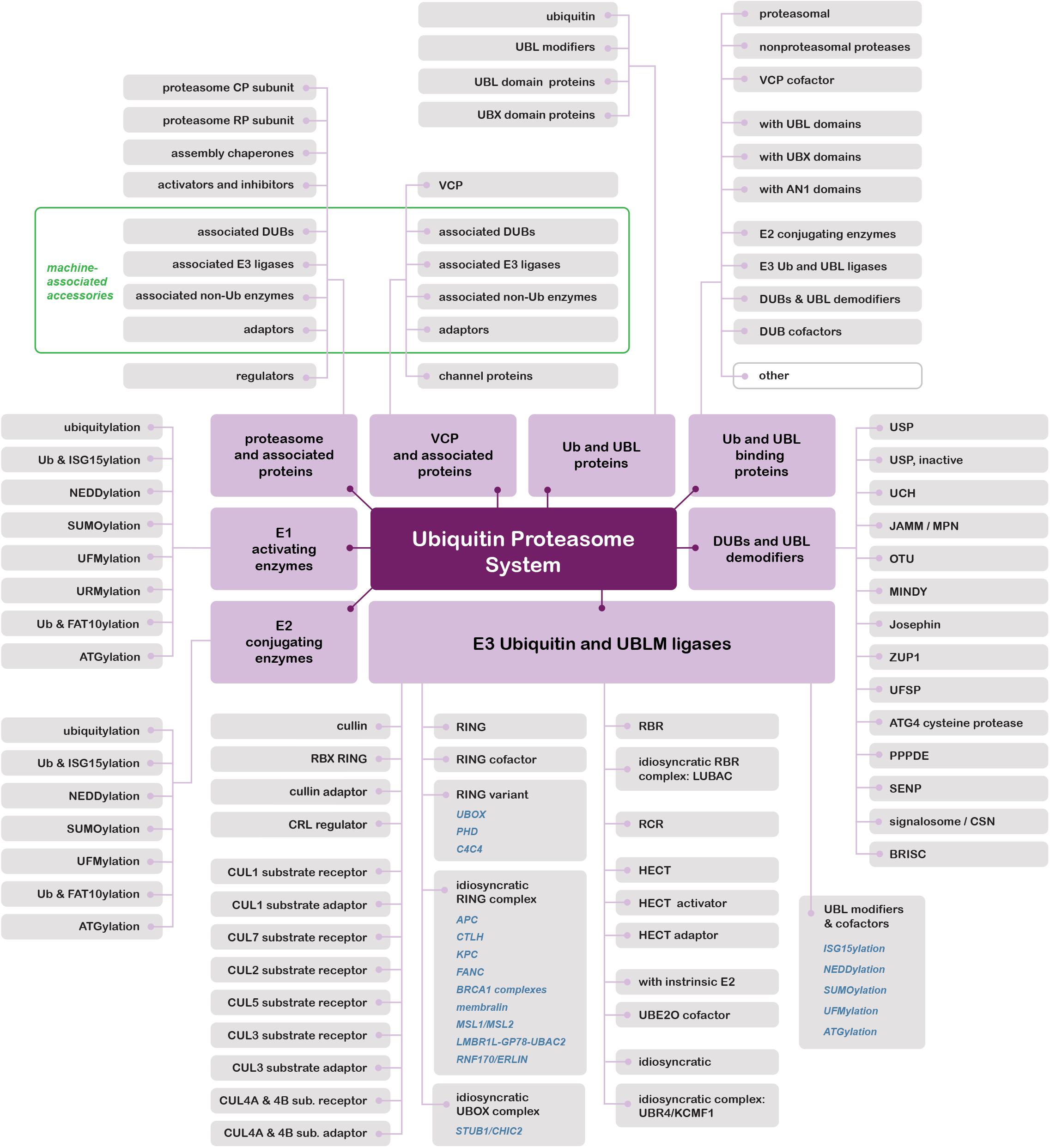
Taxonomic structure of the ubiquitin-proteasome system. The Ubiquitin-Proteasome System is shown with its Classes (fushia) and Groups (grey). For finer divisions–Type and Subtype–see **Table S1**. In a few instances, Types are shown in the figure (blue italics). The molecular machines of the UPS - the proteasome and VCP - associate with comparable suites of enzymes and cofactors, as highlighted by the green frame (upper left).

Overall, we find that within our hierarchy 1171 of 1438 UPS components fall into only one UPS Class. This demonstrates that our taxonomic scheme can organize UPS genes into functionally distinctive and human-interpretable categories that minimize ambiguity while preserving molecular context.

## RESULTS

### Approach for the survey

To generate a comprehensive accounting of the UPS (**Table S1**), we relied heavily on the modularity inherent in the system. The InterPro database^15^ has been a key reference for us in exploring the UPS as a modular system. A strength of InterPro is that it aggregates the output of independent efforts to relate protein sequence to protein function15. InterPro uses multiple designations to describe these protein sequence elements, most commonly Domains, which correspond to generally to functional or structural elements, and Families, which flag evolutionary relationships and frequently span the length of a protein. InterPro Repeats are smaller, repeated structural elements, which for the UPS most often correspond to protein-protein interaction sites. InterPro Homologous Superfamilies aggregate structurally related groups, often with low sequence homology. We identified InterPro designations, most often Domains (**Table S3**), shared by well-characterized UPS enzymes and other proteins, and used these signature domains to identify additional components. Typically, these signature domains correspond to well-understood molecular functions such as ubiquitin-directed enzymatic activities. In the course of searching for signature domains, we also identified InterPro Domains or Repeats shared by subsets of a given family. These auxiliary domains allowed subdivision of families, giving texture to what in some cases are large protein families. Some of these auxiliary domains recur in a variety of protein families, and are described as distributed domains (**Table S3**). While the majority of the proteins in our survey meet domain-based criteria for inclusion, other components of the UPS are sui generis rather than members of protein families. These are included in the UPS based on positive biochemical, cell biological, or genetic evidence, and we refer to this mode as entity-based inclusion. We used the dual approach of domain-based inclusion and entity-based inclusion in our earlier surveys as well^11,12^.

### Ubiquitin and ubiquitin-like proteins

Ubiquitin (UB), the foundational component of the UPS, shares its unitary structural motif with ubiquitin-like modifiers (UBLMs), collectively known as Type I ubiquitin-like proteins (UBLs), and with proteins bearing ubiquitin-like domains that do not participate in protein modification (Type II UBLs) (**Fig. 4**)^16^. We include here ubiquitin itself, UBLMs, and all proteins bearing structurally related domains in one Class. The mode of action of Type I UBLs can be quite different from that of ubiquitin. For example, conjugation of ATG8/LC3 (the autophagy-specific ubiquitin-like modifier) promotes protein degradation, as ubiquitin does, but only via autophagy, or the lysosome. However, ATG8/LC3 is not conjugated to proteins but to lipids, primarily phosphotidylethanolamine^27^. This reaction targets it to membranes, where it plays a key role in the genesis and trafficking of the autophagosome^28^. Ubiquitin itself is not expressed in usable form by the cell; ubiquitin precursors in the form of polyubiquitin (UBB and UBC) and ribosomal ubiquitin fusions (RPS27A and UBA52) must be processed to generate an activatable C-terminus (glycine-76). Under growth-permissive conditions, ubiquitin is expressed primarily as ribosomal fusions, yoking the production of ubiquitin to increases in ribosome abundance^29^. FAU/MNSFβ, a ubiquitin-like modifier^30^, is likewise produced as a ribosomal fusion protein and FAU itself is absent from assembled ribosomes^31^. Most ubiquitin-like modifiers must be processed prior to ligation to target proteins, resulting in a rapid release of amino acids or small peptides that has historically been considered to be of no consequence. However, deletion of the single-residue C-terminal leaving group from the polyubiquitin gene was recently shown to result in sensitivity to proteotoxic stress^32^. Another extended version of ubiquitin, known as UBB+1, is the product of frameshifting in transcription of the ubiquitin B gene, and its accumulation has been linked to neurodegeneration^33^. Among the Type II ubiquitin-like domains, the UBL and UBX domains are especially significant in that they target to the proteasome and VCP, respectively^34–36^, though it is highly unlikely that all Type II UBL domains can confer proteasome association. UBL domain proteins also have the potential to associate with any of the hundreds of proteins that have been broadly classified under likely ubiquitin binders in our survey.

**Figure 4.**
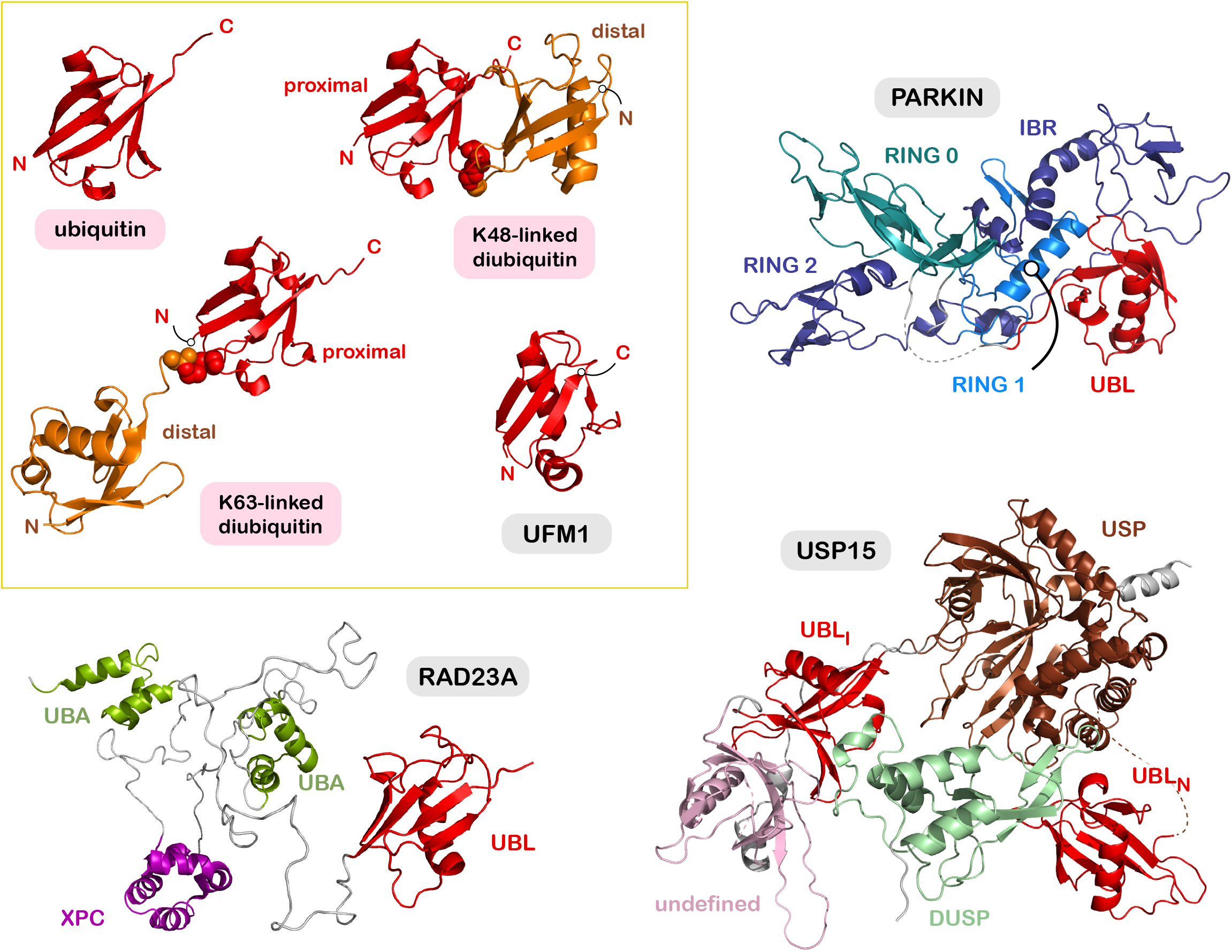
Structures of ubiquitin, UBLM, and UBL proteins. Type I UBLs are presented with in the yellow frame, and Type II UBLs outside of it. Ubiquitin and UBL domains are shown in red. For diubiquitin forms, proximal ubiquitins are shown in red and distal ubiquitins in orange. Isopeptide linkages between two ubiquitins are depicted in sphere format. Free termini for ubiquitin and UFM1 are labeled with N and C. ***Top left:*** ubiquitin and two forms of diubiquitin^17–19^. ***Center:*** ubiquitin-like modifier UFM1^20^. ***Top right:*** E3 ubiquitin ligase PARKIN (AlphaFold2 and ^21,22^). Bottom right: deubiquitinating enzyme USP15, with two UBL domains (AlphaFold2 and ^23–25^). ***Bottom left:*** proteasome shuttling factor RAD23A^26^. See **Table S4** for additional information.

While the structure of ubiquitin has been known for decades^17^, the ubiquitin and ubiquitin-like proteins are not captured by a single InterPro domain. The InterPro-designated *Ubiquitin-like domain superfamily* (IPR029071) is too broad to capture the UB/UBL proteins precisely, as it encompasses domains that are not functionally related to ubiquitin. The *Ubiquitin-like domain* (IPR000626) captures 60 family members, representing the most efficient InterPro domain for retrieving members of the UB/UBL Class, though at the same time missing more than it collects. Additional Interpro domains indicating UBLMs and UBL domains were identified by evaluating domains overlapping with IPR029071. Proteins described in the literature as containing UBL domains, but not identified as containing IPR029071, were also analyzed for the presence of InterPro and Conserved Domain Database (CDD) domains. This collection of signature domains was used to define the members of the UB/UBL Class in the main annotation (**Table S1**) and the domains themselves are presented in **Table S3**.

Not only does ubiquitin modification support protein turnover, but the attachment of several ubiquitin-like modifiers can likewise support substrate degradation by the proteasome^37–39^. Ubiquitin-like modifiers are also well known to modulate the activity of UPS pathway enzymes–most notably, cullin RING ligases are activated by the attachment of NEDD8 to the cullin subunit^40^. While the consequences of ubiquitin-like protein modification may not strictly fall within UPS processes in every case, as presently understood, they are as a group sufficiently linked in function that we chose to include all ubiquitin-like modifiers and their cognate enzymes in our survey (**Table 1, Table S1**)^37,41^.

**Table 1.**
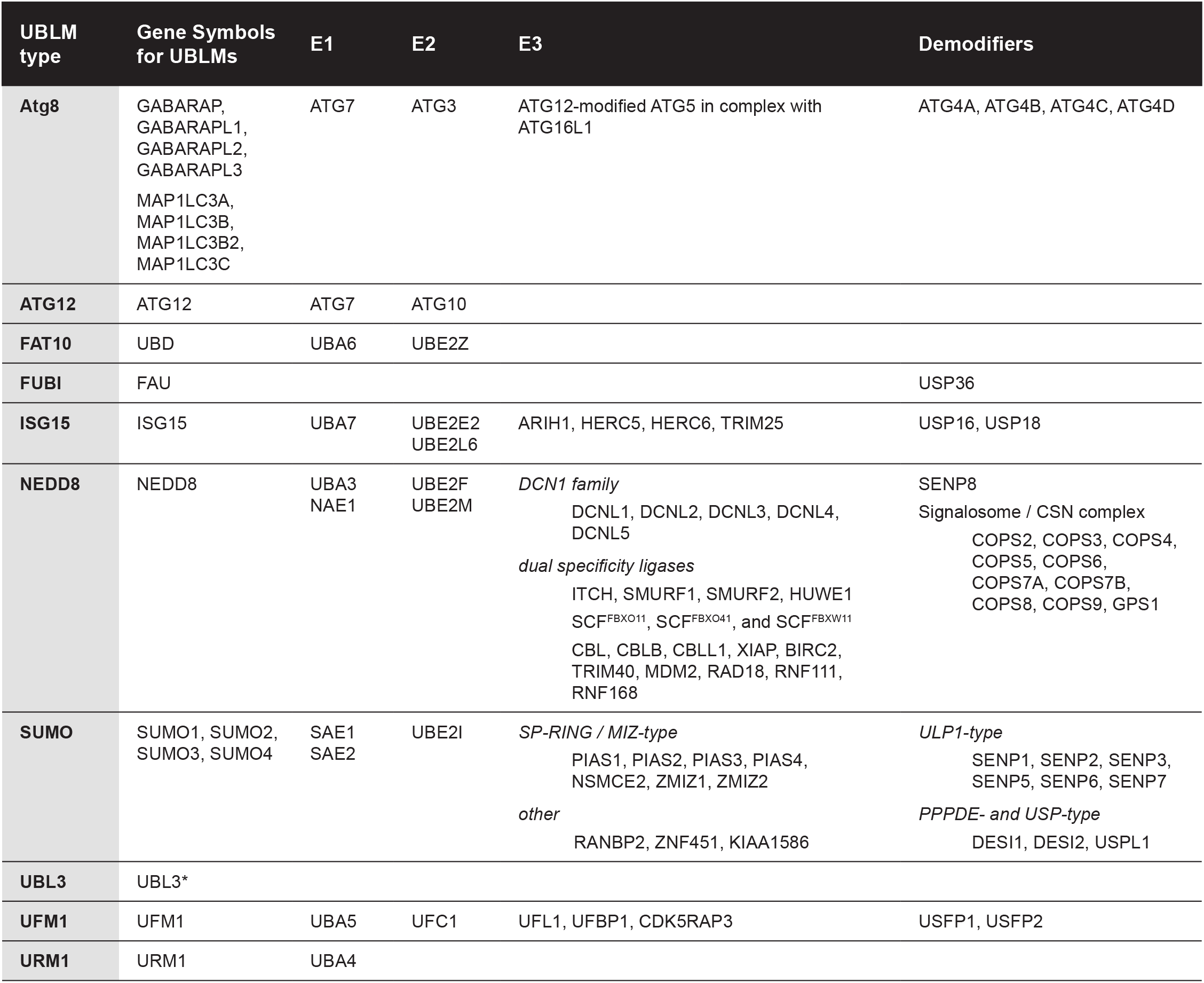
Ubiquitin-like modifiers and their cognate enzymes. *UBL3 appears to be conjugated to protein targets via a disulfide linkage^42^, rather than an isopeptide, but no responsible enzymes have been identified.

### Ubiquitin activating and conjugating enzymes

Ubiquitin enters the conjugation pathway via the action of one of two E1s, or ubiquitin activating enzymes (in humans). Ubiquitin, activated at its C-terminus by thioester formation, proceeds to undergo thiolester transfer onto one of tens of E2s, or ubiquitin conjugating enzymes, and is subsequently ligated to substrate via the action of one of hundreds of E3s, or ubiquitin ligases (**Fig. 1**). The enzyme families at the top of this pyramid are easily defined by canonical domains. While the E1 enzymes for ubiquitin and ubiquitin-like modifiers are multidomain proteins, there is one InterPro entry that unifies them: the *THIF-type NAD/ FAD binding fold* (IPR000594; **Table S3**). Most of the E2 enzymes share a single stereotypical domain (IPR000608), though the E2 enzymes for UFMylation and ATGylation are identified by other domains (IPR014806 and IPR007135, respectively). The E2 enzymes have been divided into 17 families^43^, and we have included these as Type designations in our taxonomy (**Table S1**). For the most part, the E2 Class is lightly elaborated beyond the principal domain, typically bearing what seem to be unstructured N termini, unstructured C termini, or both. These tails contribute to the specificity of E2-E3 interactions^44,45^. Two E2s, UBE2J1 and UBE2J2, include transmembrane domains placing them in the ER, two E2s, UBE2Q1 and UBE2Q2, include *RWD-like domains* (IPR006575), and one E2, UBE2K, includes a UBA domain (IPR015940). This UBA domain confers ubiquitin binding and therefore promotes ubiquitin chain formation^46,47^.

There are four catalytically inactive E2s (known as ubiquitin E2 variants or UEVs) captured by the IPR000608 domain, and these are grouped with the active E2s, as well as with the ubiquitin binding proteins. To take the UEV MMS2 as an example, it was once thought to be inactive in that it lacks a catalytic cysteine by which to form a ubiquitin thioester. Yet it is a subunit of a K63-chain forming heterodimer with UBC13 and, by orienting the acceptor ubiquitin with respect to the active site of UBC13, governs linkage specificity^48^. Perhaps the most unusual E2 enzymes are the two that function essentially as E2-E3 hybrids: UBE2O and BIRC6^49^. In fact, UBE2O also has E4 activity, in that it recognizes substrate-bound ubiquitin and preferentially ubiquitinates ubiquitinated proteins^50^. While the E2-E3 and E2-E3-E4 fusions are arguably models of efficiency, this design has barely proliferated in evolution, and probably has drawbacks such as an inherent lack of modularity.

### Properties of E3 enzymes

Once ubiquitin has moved through the tight script of E1 activation and E2 conjugation, ligation to substrate relies on a myriad cast of ubiquitin ligases. The E3s represent the most variably annotated Class, due to the breadth and complexity of this family. We apply both structural and enzymatic criteria in our basis for inclusion. For E3s to be functional as ubiquitin ligases, they must recruit an E2 charged with ubiquitin, recruit substrate, and promote the transfer of ubiquitin to substrate. The recruited substrate must also be positioned in a productive orientation with respect to the E2. Some ubiquitin ligases accomplish this with a single polypeptide, but many are composed of several subunits and when fully assembled can take the shape of arcs or rings^51–58^. CUL2^FEM1C^ serves as an example (**Fig. 5**)^59^. While only one subunit associates with the E2, and only one with substrate, the remaining three subunits are essential for the E3 activity (and are included in our survey). While E3 enzymes as a whole have been a subject of intense study for decades, many are understudied, and it is likely that some of the smaller ligases function with partners not yet identified.

**Figure 5.**
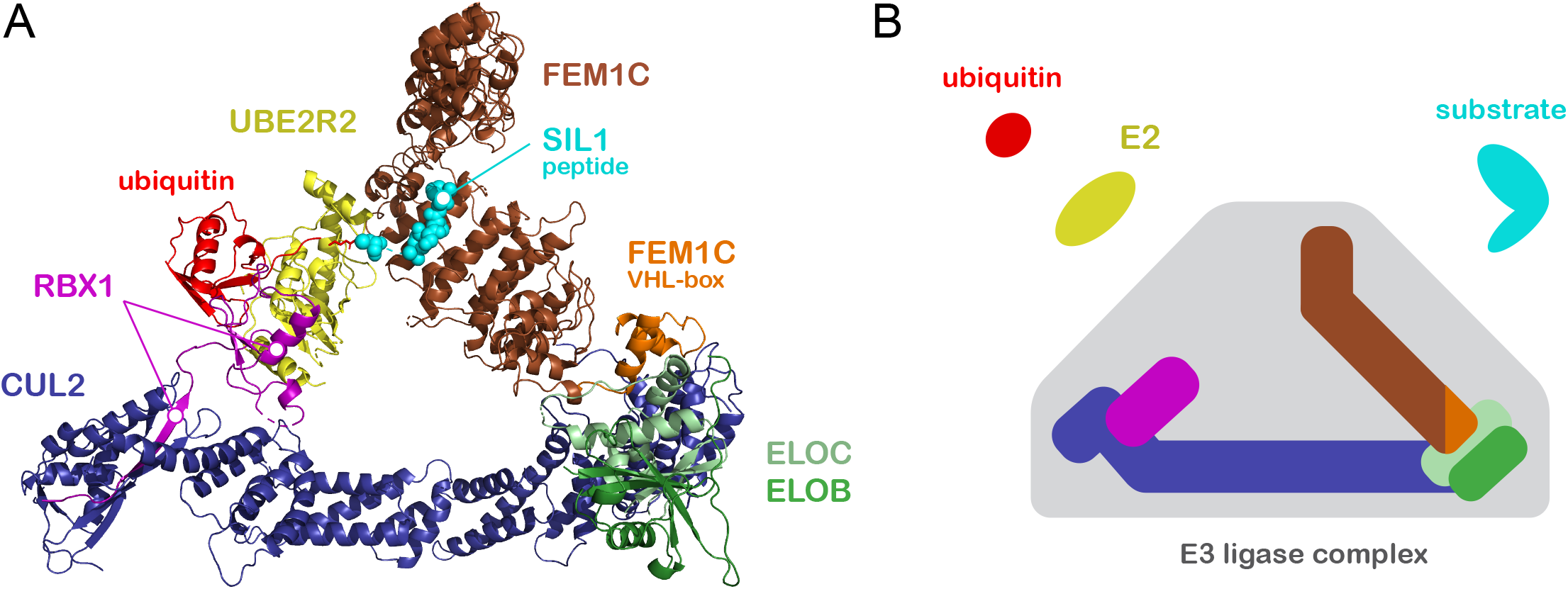
Architecture of cullin RING ligases. Structural model derived from the cryoEM map of a complex containing a cullin RING ligase bound to a ubiquitin-charged E2 poised for transfer to a substrate [PDB:8Q7R]^59^. (**A**) CUL2 (blue) with RING subunit RBX1 (magenta), cullin adaptor ELOBC (green), and substrate receptor FEM1C (brown). The adaptor-binding VHL box of FEM1C is shown in orange. UBE2R2 (yellow green), charged with ubiquitin (red), contacts the RING domain of RBX1. Substrate SIL1 (turquoise) is bound by the ankyrin repeat domain of the substrate receptor and positioned in proximity to the charged E2 for ubiquitin transfer. (**B**) A schematic of the CUL2^FEM1C^ ligase.

### E2 Recruitment by E3 enzymes

E3s are distinguished by their method of recruiting charged E2s, as well as whether they accept ubiquitin from the E2 by thioester transfer (**Table 2, Fig. 6**). The cullin RING ligases recruit charged E2s via a small RING-domain subunit specialized for this purpose—either RBX1 or RBX2. The RING-domain ligases, in contrast, use integrated RING domains to recruit charged E2s. The variant RING domains UBOX and PHD operate similarly. HECT ligases are mechanistically distinct from all of these, in that they form a thioester adduct with ubiquitin before transferring it to substrate. Their mode of E2 recognition is also distinct: charged E2s are bound by the N lobe of the HECT domain rather than through a RING. The RING-in-between-RING (RBR) ligases likewise form adducts with ubiquitin before transferring ubiquitin to substrate, though they do dock the charged E2 through an intrinsic RING domain^66^. The single member of the RING-Cys-Relay (RCR) Group^67^ similarly employs a RING domain for E2 recruitment and forms ubiquitin adducts, though notably it passes ubiquitin through two sequential thioester adducts before completing substrate ubiquitination. RZ ligases RNF213 and ZNFX1 recruit distinct E2s through dissimilar domains, though they each form thioester adducts with ubiquitin via their RZ domains^68–71^.

**Table 2:** E2 recognition components of E3 ligases. *The 19 gene products not reflected in this number include RBX1 and RBX2, MARCH ligases (4 of 11), UBOX5, RBR ligases (10 of 14), RCR ligase MYCBP2, and RZ ligase RNF213.

| E3 ligase group | Related Interpro Domain | Occurrences in survey | Mode of E2 docking | Transthiolation by E3 |
| --- | --- | --- | --- | --- |
| **Cullin RING** | IPR016158 | 9 | via associated RBX1 or RBX2 (RINGS) | no |
| **RING** |  |  | via RING domain | no |
| most groups, complexes | IPR001841 | 281 |  |  |
| MARCH | IPR011016 | 11 |  |  |
| Pellino | IPR048335 | 3 |  |  |
| IRF2 binding | IPR044882 | 3 |  |  |
| CTLH complex | IPR044063 | 3 |  |  |
| FANCL complex | IPR026850 | 1 |  |  |
| **RING variant** |  |  | via variant RING domain | no |
| UBOX | IPR003613 | 9 |  |  |
| select PHDs | IPR001965 | 6 |  |  |
| C4C4 (TMEM129) | none | 1 |  |  |
| **HECT** | IPR000569 | 28 | via N-lobe of HECT domain | yes |
| **RBR** | IPR044066 | 14 | via RING1 of TRIAD domain | yes |
| **RCR** | none | 1 | via RING and loop extension | yes (twice) |
| **UBR4** | IPR025704 | 1 | via hemiring-UZI module | no |
| **RZ** | IPR046439 | 2 | variable | yes |
| **E2-E3 fusions** | none | 2 | intrinsic E2 | no |
| **idiosyncratic** | none | 13 | variably known | variably known |

**Figure 6.**
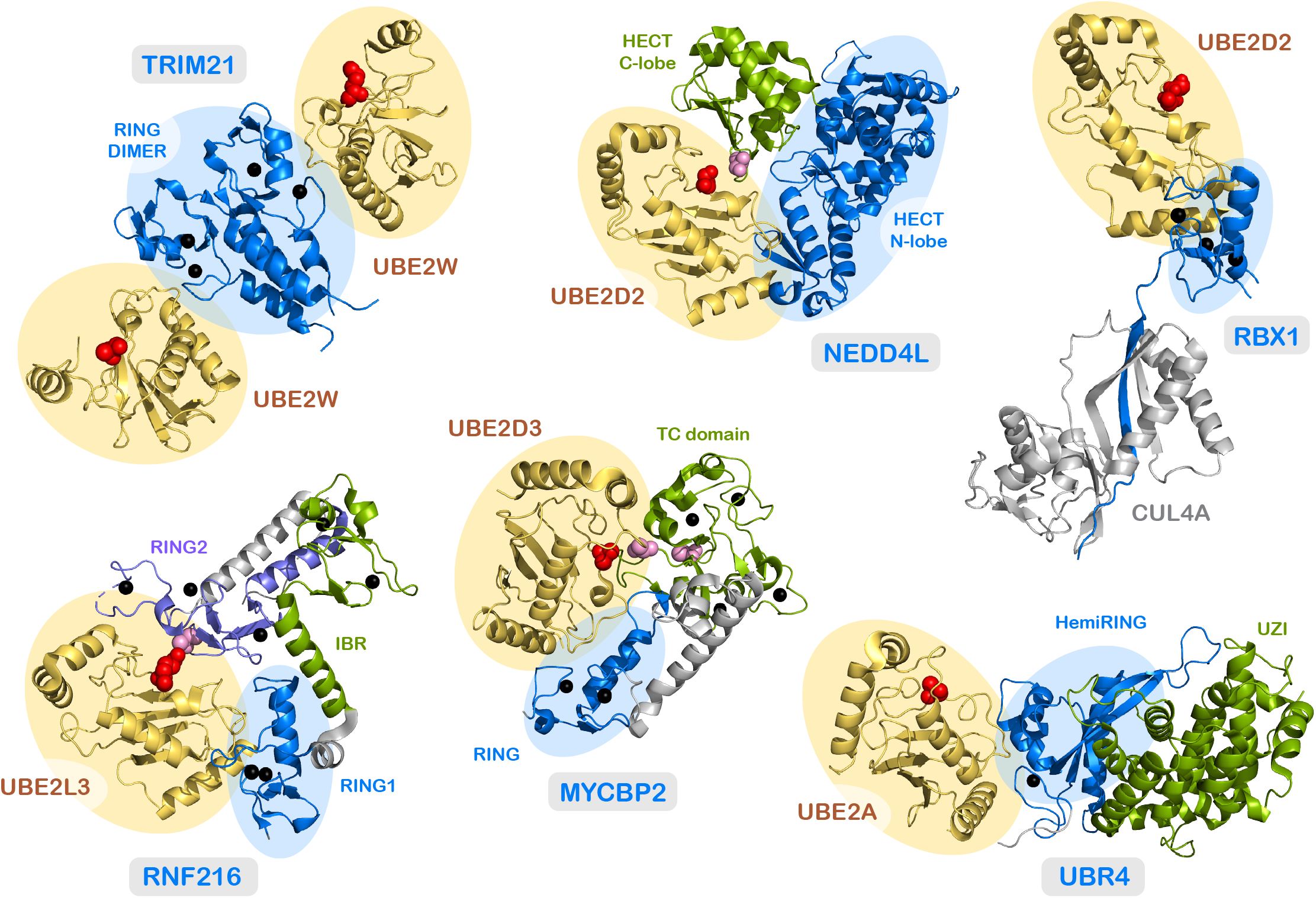
Varieties of E2 engagement. E3s engaging cognate E2s using a variety of motifs are shown. E2s are shadowed in yellow and their active site cysteines are shown in red as a space filling model. Portions of E3s that dock E2s are shadowed in blue. Catalytic cysteines for E3s are depicted in magenta. Zinc ions are rendered as black spheres. Ubiquitin, when present in structures, was omitted for clarity. **Clockwise from top left:** TRIM21 engages UBE2W via a dimerized RING domain60; HECT ligase NEDD4L engages UBE2D2 via the N-lobe of the HECT domain61; RBX1 docks into cullin CUL4A by completing a beta sheet in the RBX1 binding domain, then recruits UBE2D2 via its RING domain62; UBR4 contacts UBE2A via a unique hemiRING domain that is embedded in an UZI domain63; RCR ligase MYCBP2 engages UBE2D3 via a RING domain which supports transfer of ubiquitin sequentially to two cysteines in the TC domain64; and RBR ligase RNF216 docks UBE2L3 via one RING domain, which supports the formation of a covalent adduct with the second RING domain65. See Table S4 for additional information.

### Specificity of E2-E3 pairings

Given that E3s vastly outnumber E2s, at least some E2s must work in concert with many E3 partners. However, insofar as the mapping function is understood, it appears to be most typically a many-to-many map rather than one-to-many. Some fairly specific E2-E3 pairs are known^45,72^, but they may be atypical. A comprehensive determination of E2-E3 pairings has not been achieved, though many robust biochemical pairings are known^44^. In the course of the Predictomes project^73^, about 16% of the 15,000 possible pairings were modeled with AlphaFold Multimer, generating ipTM scores. (These scores reflect the overall confidence with which AlphaFold positions proteins with respect to each other.) This project used machine learning to calibrate AlphaFold Multimer predictions in light of experimental evidence, and the resulting SPOC (Structure Prediction and Omics-informed Classifier) scores reflect this new weighting.

**Table S5** lists all E3 components expected to interact with E2s, and both the ipTM and SPOC scores as available. Of the 349 E3 components expected to associate with E2s and also examined via AlphaFold Multimer, roughly half were modeled at high confidence with one or more E2s. In contrast, in the BioPlex 3.0 database, which is compiled from proteome-wide affinity screens, relatively few E2-E3 pairs were recovered. For example, while UBE2D2 shows high-confidence modelling with 126 E3s, only three E3s were identified as BioPlex hits. Similarly, for UBE2L3, high confidence models were obtained with 72 E3s, but only four were recovered in BioPlex 3.0. The transient nature of the E2-E3 interaction presumably hampers the identification of authentic E2-E3 pairings via affinity screens. Beyond the issue of affinity, a number of valid interactions may have been missed because the relevant proteins were not expressed under the experimental conditions of BioPlex.

### Ubiquitin ligase families and their components

#### Cullin RING ligases

Cullin RING ligases (CRLs) constitute one of the two largest families of ubiquitin ligases^8,74–76^, the other being the RING ligases (**Table S1, Table S2**). The cullin subunits serve as scaffolds to nucleate multicomponent E3s. Cullins share a variety of overlapping InterPro domains (**Table S3, Fig. 7A**). The *Cullin-like, alpha+beta domain* (IPR059120) binds the small RBX1-type RING domain proteins^77^, which in turn recruit E2 enzymes. The C-terminal *Cullin, neddylation domain* (IPR019559) is modified by NEDD8, a requirement for CRL activation^78^. The *Cullin, N-terminal domain* (IPR001373) is a superhelical domain comprising roughly 20 alpha helices and forms the backbone of the scaffold. The N-terminal four-helix bundle of this domain recruits adaptors which in turn recruit substrate receptors (**Fig. 7B and 7C**). The metazoan cullins CUL7 and CUL9 lack this four-helix bundle and engage substrates in a manner distinct from that of the canonical cullins. ANAPC2, a component of the Anaphase Promoting Complex (APC), lacks the neddylation domain, and is not considered to be a cullin but a degenerate offshoot of the family. CACUL1 lacks both the neddylation and RING-binding domains and is not expected to support E3 activity. It has been found to stabilize the oxidative-stress-sensitive transcription factor NRF2, probably by suppressing the CUL3^KEAP1^ ligase^83^, and is retained in the survey and filed with the cullins.

**Figure 7.**
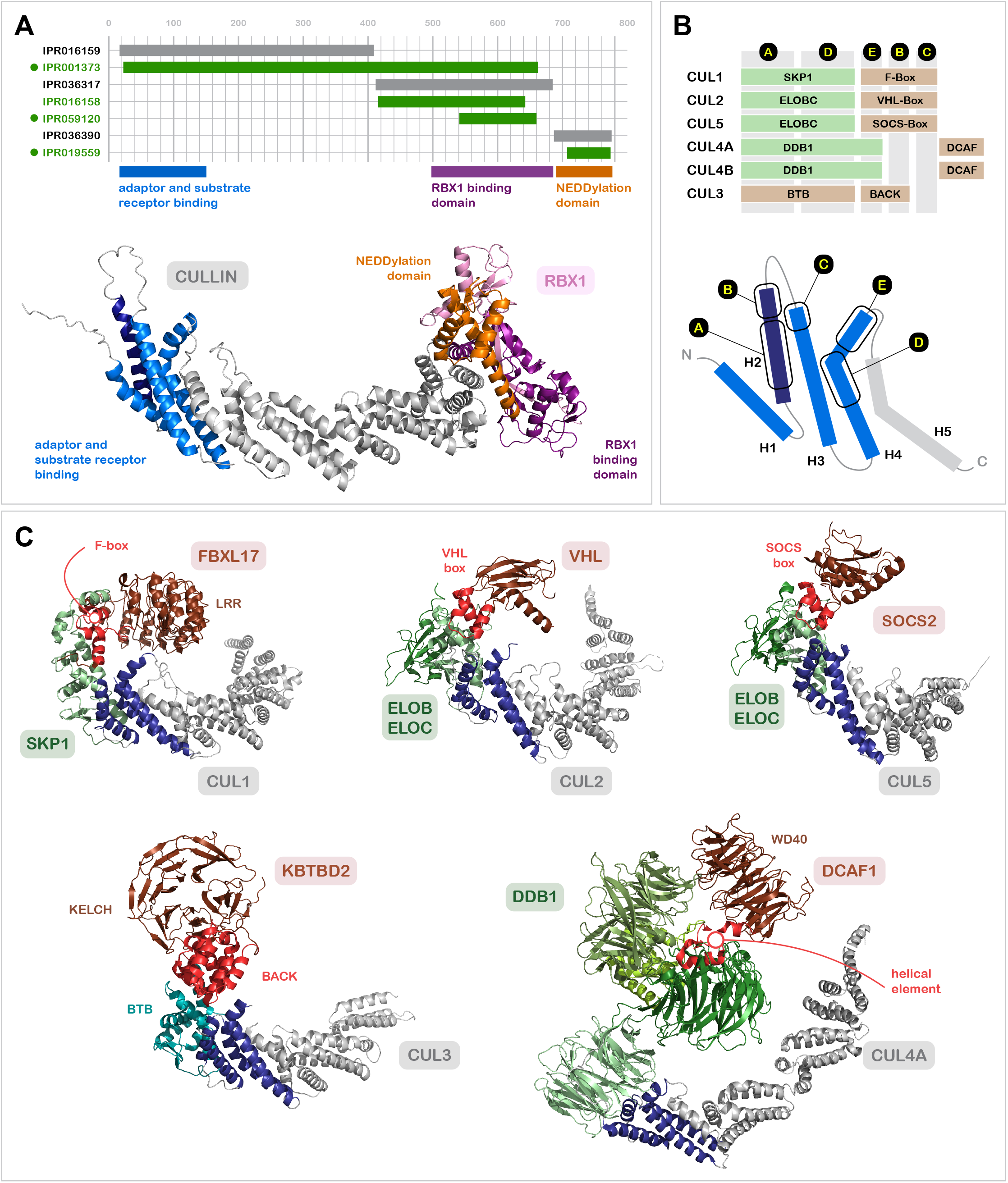
Cullin determinants engaging the cullin adaptors and substrate receptors. (**A**) Structural features of cullins. ***Top:*** Linear map of a generic cullin, with grey and green bars depicting InterPro homologous families and domains, respectively. Dots indicate domains referenced in the main text. Shared features of CUL1, CUL2, CUL3, CUL4A, CUL4B, and CUL5 are shown in colored bars below the diagram. ***Bottom:*** shared features mapped onto an AlphaFold3 model of CUL1 in complex with RBX1. (**B**) Comparison of adaptor and substrate receptor docking by cullins. ***Bottom:*** a cartoon of a portion of a generic cullin is shown with binding determinants highlighted by letters A-E. ***Top:*** a schematic representing engagement of cullin subunits with their cognate adaptors (mint green) and substrate receptors (light brown). CUL1, CUL2, and CUL5 engage adaptors through elements A and D. Substrate receptors associate principally with adaptors, but also form contacts with elements B, C, and E of the cullins. CUL4A and CUL4B use a large adaptor, which holds receptors (DCAFs) at a considerable distance from the cullins. CUL3 does not have a dedicated adaptor such as SKP1, but associates with substrate receptors that have an adaptor-like BTB domain. Like true adaptors, this adaptor-like domain engages the A and D elements of the cullins. The adjacent BACK domain associates with the B and E elements used by other substrate receptors. (**C**) Canonical cullins in complex with their adaptors and representative substrate receptors. Cullin domains are depicted (without RBX1-binding domains or NEDDylation domains) in grey, with the four-helix bundle that binds adaptors and substrate receptors shown in blue. Adaptors for CUL1^79^, CUL2^59^, CUL5^80^, and CUL4A^81^, and are shown in shades of green, with the portions that contact receptors shown in light green. For the substrate receptors, adaptor and cullin binding portions shown in red and the presumptive substrate binding domains shown in brown. In the CUL3 assembly^82^, the BTB portion of receptor KBTBD2 is shown in teal, and the BACK domain in red. See Table S4 for additional information.

The cullin family is ancient, and members have been suggested to map to three hypothetical ancestral genes: Culα, Culβ, and Culγ (**Table 3**)^84^. The grouping of CUL1, CUL2, and CUL5 correspond to Culα, with CUL3 and CUL4 corresponding to Culβ and Culγ, respectively. The mode of substrate recognition used by each of these cullin groups reflects, to a degree, their ancient origin. CUL1 associates with cullin adaptor SKP1, and this complex recruits substrate receptors through the F-box domain. Proteins bearing this motif are characterized by the *F-box domain* (IPR001810) and *F-box-like domain superfamily* (IPR036047), with neither alone found in all F-box proteins (**Table S3**). The F-box proteins also contain various domains that are known to interact with substrate, the most common being leucine-rich repeats (IPR032675) and *WD40 repeats* (IPR001680). These are broadly-used protein-protein interaction domains. The *F-box associated (FBA) domain* (IPR007397) is specific to the CUL1 substrate receptors, though it does appear in NCCRP1/ FBXO50, which lacks a canonical F-box.

**Table 3:**
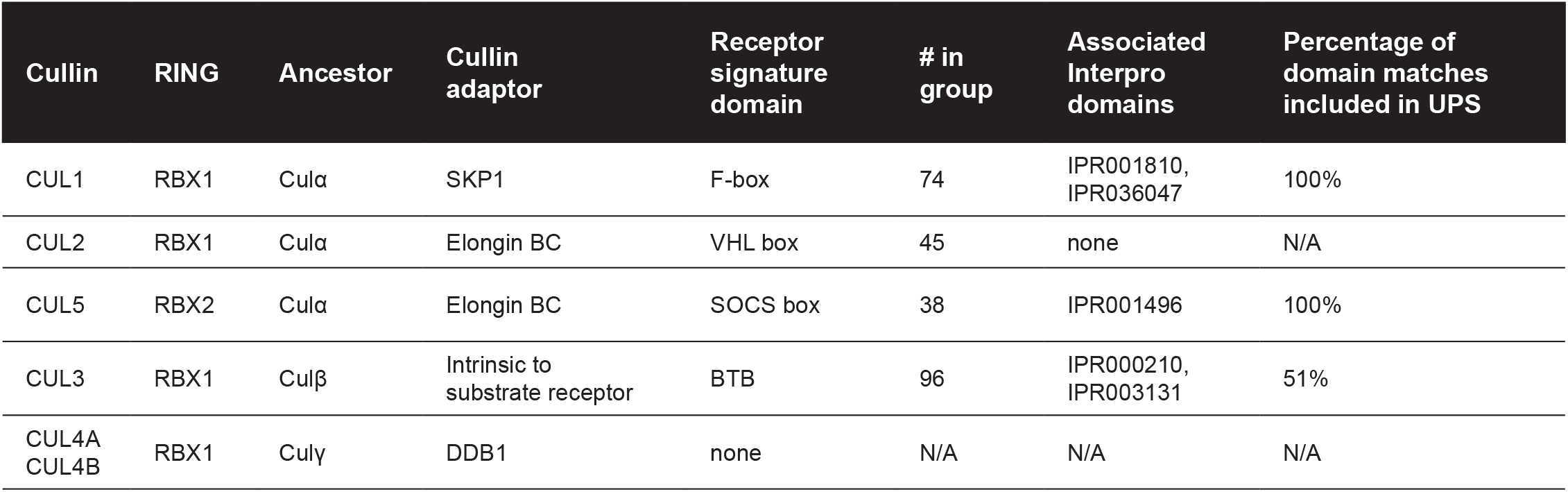
Composition of the modular Cullin Ring Ligases.

| Cullin | RING | Ancestor | Cullin adaptor | Receptor signature domain | # in group | Associated Interpro domains | Percentage of domain matches included in UPS |
| --- | --- | --- | --- | --- | --- | --- | --- |
| CUL1 | RBX1 | Culα | SKP1 | F-box | 74 | IPR001810, IPR036047 | 100% |
| CUL2 | RBX1 | Culα | Elongin BC | VHL box | 45 | none | N/A |
| CUL5 | RBX2 | Culα | Elongin BC | SOCS box | 38 | IPR001496 | 100% |
| CUL3 | RBX1 | Culβ | Intrinsic to substrate receptor | BTB | 96 | IPR000210, IPR003131 | 51% |
| CUL4A<br>CUL4B | RBX1 | Culγ | DDB1 | none | N/A | N/A | N/A |

Both CUL2 and CUL5 recruit their substrate receptors using the heteromeric adaptor ElonginBC^85,86^. Despite employing the same cullin adaptor complex, these two cullins have distinct sets of substrate receptors. CUL2 recruits substrate receptors through VHL boxes, and CUL5 through SOCS boxes. Each of these domains is divided into a BC-box, which contacts ElonginBC, and cullin-box, which distinguishes between the two cullins^87,88^. The ***SOCS box domain* (IPR001496)** efficiently identifies the CUL5 substrate receptors, though we propose seven SOCS box proteins that do not have the canonical motif (**Table S1**). CUL5 substrate receptors most commonly use *Ankyrin repeats* (IPR002110) and *SH2 domains* (IPR000980) for substrate recruitment, which are of course broadly represented in the proteome. Despite the similarities between CUL5 and CUL2, the substrate receptors of CUL2 are not easily identified using a canonical domain. The CUL2 substrate receptor VHL interacts with CUL2 via the VHL box (IPR024048), but other proposed CUL2 receptors do not share this domain, and in fact have no common signature domain. The CUL2 receptors use a variety of substrate interaction domains, with leucine-rich repeats (IPR032675) being dominant. Just as CUL2 and CUL5 share the adaptor ElonginBC, CUL4A and CUL4B share DDB1^89–91^.

The central beta propeller of DDB1 contacts the N-terminal four-helical bundle of CUL4A and CUL4B, and the flanking beta propellers form a deep cleft for the binding of the substrate receptors. While SKP1 and ElonginBC are modest in size, and allow the substrate receptors to simultaneously dock with the cullin and cognate adaptor, DDB1 is quite large, interposing at least 40Å between cullin and receptor (**Fig. 7C**). In some cases, the cell exploits the size of this adaptor. CUL4A substrate receptor DDB2 docks into DDB1 via a helical hairpin and one surface of its WD40 domain. Basic residues on the opposite face of the WD40 domain contact UV-damaged DNA, and this large complex sweeps out a “ubiquitination zone” in proximity to DNA photodamage^92^. Putative CUL4A/CUL4B substrate receptors were identified in four proteomic studies^90,91,93,94^, and were found to be substantially enriched in WD40 repeats (see Methods). Subsequent studies showed that while the WD40 domains are proximal to DDB1, the most intimate contacts to DDB1 reside in alpha helical elements that are adjacent to the WD40 domains and insert into the DDB1 cleft (**Fig. 8**). The alpha helical element is likewise a binding determinant for CUL4A/CUL4B substrate receptors lacking a WD40 repeat (**Fig. 8**). We classified as CUL4A/CUL4B substrate receptors proteins that were identified by DDB1-affinity purification and that also modeled well with DDB1 (**Fig. S1**). We also included homologs of strong candidates, as well as proteins with poor DDB1 modelling that are nonetheless supported by biochemical evidence as promoting CRL4 ligase activity. We excluded as CUL4 substrate receptors proteins from the initial studies that lacked any of the support described above^93,94^.

**Figure 8.**
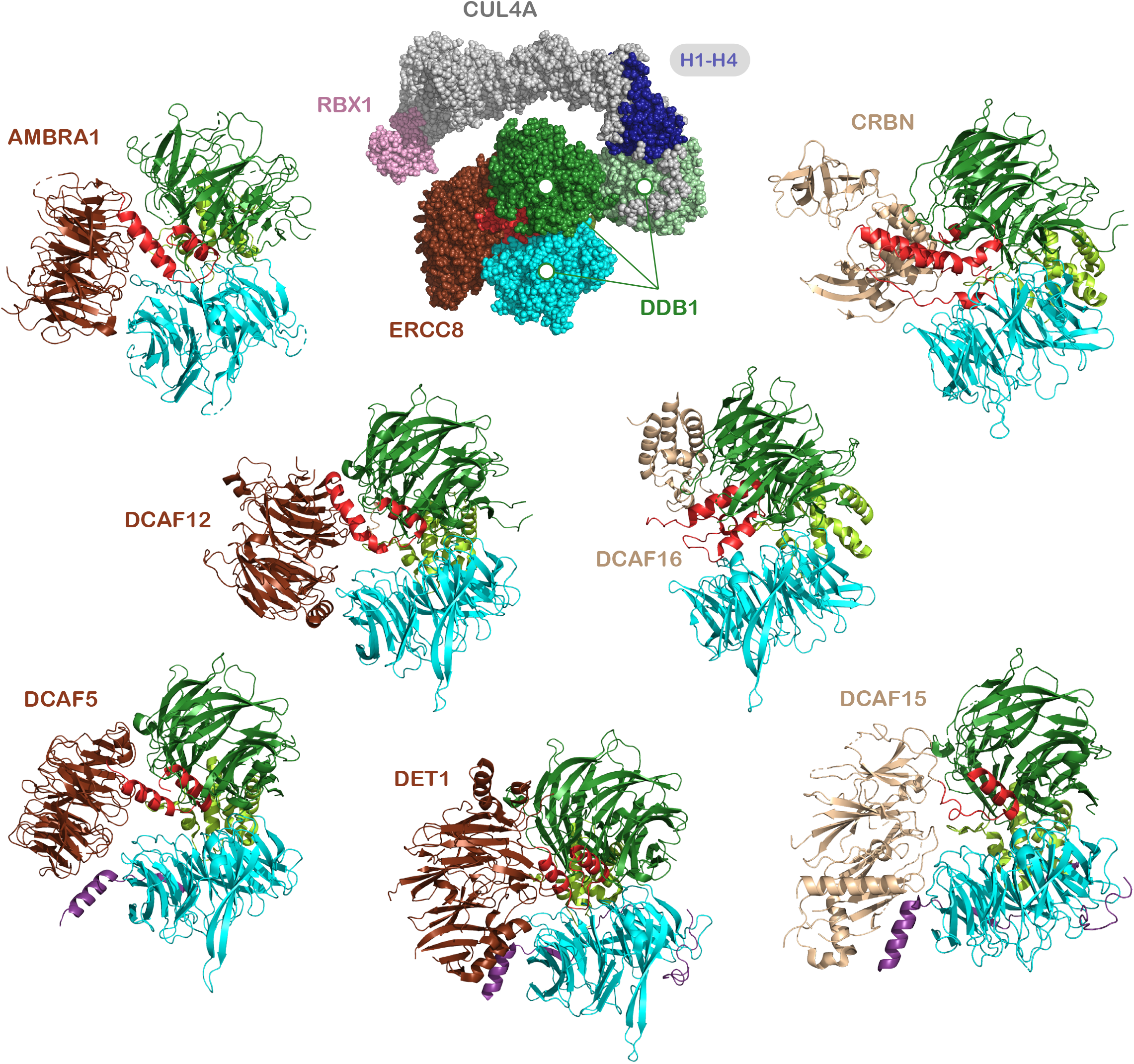
CUL4 substrate adaptors. ***Top center:*** CUL4 substrate receptor ERCC8 (brown and red) solved with DDB1 (sage, green, and teal), CUL4A (dark blue and grey), and RBX1 (pink), shown in spacefill^95^. The remaining structures only contain DDB1 and the CUL4 substrate receptors. ***Left and bottom center:*** CUL4 substrate receptors with WD40 domains (brown)^96–99^. ***Right:*** CUL4 substrate receptors without WD40 domains (beige)^100–102^. For all structures, helices in red are the components of CUL4 adaptors that form the majority of contacts between the CUL4 substrate receptors and the flanking beta propellor domains of DDB1. See **Table S4** for additional information.

CUL3 recruits substrate receptors not through an intermediate cullin adaptor, but through an adaptor domain intrinsic to CUL3 substrate receptors^103–106^, namely the BTB domain (IPR000210). This binds the same regions of the cullin four-helix bundle that are recognized by the adaptors for CUL1, CUL2, and CUL5 (**Fig. 7B**), and is found within the *SKP1/ BTB/POZ domain superfamily* (IPR011333), which also includes SKP1 and ElonginBC. The BTB domain (IPR000210) is found in 178 human proteins, but we consider only half of these to be CUL3 receptors, as BTB domains are not necessarily sufficient for productive binding with CUL3^107–109^. Valid CUL3 receptors mostly partition into two groups (**Fig. S2**). The larger group is characterized by a BACK domain (IPR011705) appended to the BTB domain, with the 3-box element of the BACK domain mimicking the contacts formed between CUL1, CUL2, and CUL5 and their substrate receptors (**Fig. 7B and 7C**). The BACK domains are not exhaustively captured by IPR011705; we identified an additional 20 BACK domains by evaluating each BTB protein for an adjacent BACK domain (see **Table S3**). KELCH repeats (IPR006652) are found in two-thirds of the BTB-BACK-type CUL3 receptors. All BTB-BACK proteins were modeled with CUL3 using AlphaFold3, and those with weaker scores are included in our survey and flagged as variants.

The second group of CUL3 receptors carries the KCTD-type BTB domains (IPR003131). In total, there are fifty KCTD-type BTB domain proteins, partitioning into Type I and Type II. The Type I group was recently evaluated systematically via biochemistry and modelling^110^, and only two-thirds were found to associate with CUL3. We adopted those findings wholesale in our survey. Note that the KCTD-type BTB proteins that host CUL3 tend to assemble into pentameric forms with CUL3 subunits emanating from BTB rings^111^ so as to contact not one but two faces of the BTB domains^110^. The Type II group carries *Ion transport domains* (IPR005821) adjacent to the BTB domain, and in at least some cases steric hindrance should prevent association of these proteins with CUL3. The last large group of BTB proteins, which carry between two and fourteen C2H2-type zinc fingers (IPR013087) at their C-termini, generally lack key features for driving CUL3 engagement. Most members of this set are described in UniProt as transcriptional regulators. Until recently, CUL3 substrate receptors were thought to consist entirely of BTB domain proteins, but recent work has shown that ZSWIM8 complexed with ElonginBC forms a functional complex with CUL3, potentiated by a specialized sequence between the BC-box and the CUL2-box^58^.

### RING ligases

While the cullin RING ligases recruit E2s by means of an adaptor subunit, the remaining Groups of ligases recruit charged E2s through intrinsic domains. The RING ligases constitute the largest of these Groups, numbering ∼300^112–114^. The RING ligases in our survey are extremely varied, ranging for example in length from about 150 residues to more than 5000. As indicated in **Table 2**, more than 90% of the RING ligases share the RING-type zinc finger (IPR001841) domain. Within the RING ligase Group, a few Types of RING ligases are designated by other signature domains; the three Pellino ligases share IPR048335, the eleven MARCH ligases share IPR011016, the three proposed IRF2BP ligases share IPR044882. The UBR RING ligases do not share an InterPro domain that corresponds to RING activity, but they have been characterized experimentally^115^ and are annotated with RING domains from the CDD. In a few cases, comparison to paralogs allowed the identification of RING domains (RFPL2, PEX12, and VPS1), and these are also annotated with CDD domains. The RING ligase family has been sorted into more than 30 Types with homologous regions beyond the RING domain^114^. We largely adopt these categories, and introduce a few new ones based on shared properties: RING ligases with transmembrane domains, with ubiquitin or UBL binding domains, or bearing additional enzymatic functions.

The RING ligase family contains one exceptionally large family, known as the TRIM ligases. Numbering more than 80, these were parsed previously into 12 subsets^116,117^, and we follow that organization as the basis for Type designations for the TRIMs. More than half of the TRIMs contain a PRY-SPRY substrate-recognition domain (IPR001870, TRIM Class IV), but this domain is not exclusive to TRIMs or to RING ligases. Structurally, TRIMs are notable in that their RING domains are separated from their substrate interaction domains by a single alpha helix of typically 100 residues or more, in most cases containing a break in the helix toward its C-terminal end, which allows for the formation of a hairpin structure. These long helices are typically involved in homodimer or heterodimer formation with family members, and some higher order assemblies are known^117–119^. Dimerization places the RING domains in proximity to each other, which promotes the release of ubiquitin from the E2 and transfer of ubiquitin to substrate^120^. Recently, the TRIMs were systematically evaluated for homodimer formation to activate the TRIMs^117^. Some were found to be inactive, at least in these assays, calling into question whether they are true E3s. Nonetheless, we include all of the TRIMs in our survey. RINGless TRIMs, although not expected to recruit E2s, were included in our survey because they may modulate TRIM activity. Interestingly, while we assembled this survey, we noticed that 11 TRIMs are modeled in AlphaFold2 as having folded central helices, which may modulate dimerization (**Table S1**).

### Idiosyncratic RING ligase complexes

Of the myriad RING ligases, many function in the context of larger complexes. Annotation efforts typically include only the catalytic portion of these complexes; we include all subunits. While we reference these ligases as idiosyncratic, they are no less interesting for their individuality. CTLH, conserved through eukaryotes, is built around a RING heterodimer and at least three times as many noncatalytic components. The core complex assembles into circular supramolecular complexes of 1 MDa or more, which are capable of ubiquitinating oligomeric structures through avid interactions^121^. The anaphase promoting complex (APC) is built around a catalytic core containing one RING subunit and one degenerate cullin, but this core is embedded in a much larger assembly. Using exchangeable substrate receptors, the APC drives the cell cycle by promoting timed degradation of cyclins, securin, and tens of other proteins. The much smaller KPC complex also controls cell cycle progression, driving the G_0_ to G_1_ transition. It is composed of the RNF123 RING subunit, which contains a SPRY domain, and a noncatalytic UBAC subunit. The latter encompasses two UBA domains, which bind the ubiquitin moieties of modified substrates, and a UBL domain, which promotes targeting of substrate to the proteasome.

Several RING complexes are known to engage damaged DNA, and failures of these complexes result in cancers and anemias. The first is the FANC core complex, which promotes the repair of interstrand DNA crosslinks. Mutations in FANC complex components give rise to Fanconi Anemia^122^. The FANC core complex is composed of one RING domain protein, which also contains a ubiquitin binding element, and an additional six noncatalytic subunits. The ubiquitin binding domain of FANCL supports monoubiquitination of pathway-critical FANCD2 and FANCI^123^. The repair of DNA double-strand breaks (DSBs) involves multiple E3 enzymes that target histone H2A for modification in the neighborhood of the DSB, including RNF8, RNF168, and, downstream of them, BRCA1-BARD1^124^. Each subunit of BRCA1-BARD1 contains a RING domain, and the complex dimerizes through helical bundles flanking the RING domain. It may be the case that only recruitment of the E2 through BRCA1 is required for full activity^125^, though BRCA1 is strongly activated by BARD1. DNA repair is also promoted by ubiquitination of histone H2B, notably by the Bre1 ligase complex^126–129^. In addition to resolving DNA damage, E3s engage DNA in modulating transcription. The RING ligase MSL2 ubiquitinates histone H2B as well, and this activity is strongly activated by forming a heterodimer with MSL1^130^. Note that this heterodimer is embedded in a larger histone acetylation complex^131^.

Misfolded proteins present in the ER are degraded via the ERAD pathway (ER-associated degradation) in which substrates are exported back to the cytosol–a process known as retrotranslocation– before the proteasome can degrade them. In the canonical pathway of retrotranslocation, a ubiquitin ligase, Hrd1, constitutes the primary transmembrane channel in yeast, ubiquitinating the substrate once it is exposed to the cytoplasm^132^. Extraction from the membrane is then driven by VCP^133^, which recognizes the ubiquitin groups added by Hrd1. In humans, a variety of ubiquitin ligase complexes are embedded in the ER membrane^134^. To cite one example, the active component of the membralin complex is the RING ligase RNF185, a small protein consisting primarily of a transmembrane hairpin anchoring a RING domain on the cytosolic side of the ER membrane. RNF185 functions in complex with three other membrane proteins, including the much larger TMEM159, which may be involved in substrate recognition in the ER lumen, and TMUB1 and TMUB2, both of which have UBL domains on the cytosolic side. Another includes the LMBR1L-GP78-UBAC2 complex, which includes the RING ligase GP78/AMFR. The N-terminal portion of AMFR is embedded in the ER membrane, and the C-terminal half projects into the cytoplasm. This portion engages UBE2G2, its cognate E2, not only through a RING domain but also a second domain which promotes high affinity binding. Connecting these two is a CUE domain, which associates with ubiquitin. AMFR associates with two other membrane proteins, UBAC2 and LMBR1L, the former including a UBA-type ubiquitin binding domain and the latter engaging with components of the WNT signaling pathway.

### Variant RING ligases

In the RING domain of the RING ligases, two zinc ions coordinate a combination of eight cysteines and histidines to form a cross-brace structure which forms the basis for the RING signature. The 40-60 residue motif contains a small three-strand beta sheet and an alpha helix. Loops connecting these elements provide docking sites for E2 enzymes. To dock the E2s, the *U-box domain* (IPR003613) mimics this arrangement through a collection of hydrogen bonds rather than zinc coordination^135^. As compared with true RING domains, the collection of UBOX ligases is small, numbering only eight, and we put them in a distinct Group. There are two proteins, RBBP6 and TRIM21, that have overlapping RING (IPR001841) and UBOX (IPR003613) motifs, and we place these in the RING rather than UBOX Group. One protein, UBOX5, has distinct UBOX and RING domains, and it is the UBOX that is required for E2 association^136^.

Like the RING domain, the PHD domain (IPR019787, IPR001965) coordinates two zinc ions in a cross-brace configuration^137^. By and large, PHD domains associate with methylated histones rather than E2s^138^, but six PHD domain proteins have been found to function as ubiquitin ligases. These include AIRE, G2E3, ING4, JADE2, KMT2A, and KMT2B^139–144^. While lacking a canonical PHD domain, UBR7 bears a PHD-like domain that supports its ubiquitin ligase activity, and it is included as a variant PHD^145^. PHD and RING domains occasionally occur in the same protein. Six RING ligases have predicted PHD domains entirely coincident with predicted RING domains (BAZ1B, NFX1, NSD2, PHRF1, UHRF1, and UHRF2). In contrast, KMT2C and KMT2D have multiple predicted PHD domains that do not fully overlap with predicted RING domains—we categorize these as canonical RING ligases. Five RING ligases have PHD domains that are clearly distinct from their RING domains, and these include SHPRH, PHF7, and the three Class VI TRIMs (TRIM24, TRIM28, and TRIM33).

### HECT ligases

Unlike RING ligases, the HECT ligases form thioester adducts with ubiquitin, a property of the HECT domain^146^. This domain is divided structurally into N-terminal and C-terminal lobes connected by a flexible hinge. E2s associate via the N-terminal lobe and donate ubiquitin to a cysteine in the C-terminal lobe. Ubiquitin is transferred from this cysteine to the substrate. The HECT ligases constitute a Group of 28 proteins which are defined by a single domain (*HECT domain*, IPR000569). HECT ligases are elaborated by a variety of domains which determine substrate specificity and cellular context. The most common substrate interaction motifs include WW (IPR001202) and RCC1 (IPR009091). The smallest of the HECT ligases are roughly 80 kD, and several weigh in around 500 kD: HERC1, HERC2, and HUWE1. The latter adopts a ring-shaped architecture decorated with protein interaction modules, including ubiquitin binding domains^53^. The ubiquitin binding elements provide a means of recruiting soluble substrates and aggregates that carry a high density of ubiquitin chains, and promoting amplification of the ubiquitin signal^147^.

### RBR, RCR, and RZ ligases

The RBR and RCR ubiquitin ligases combine features of RING and HECT ligases in that they recruit charged E2s via RING or RING-like domains, however they form thioester adducts with ubiquitin before transferring it to substrate^66,67^. For the 14 RBR ligases, the functional trio of an E2-binding RING1 domain, an IBR domain, and a catalytic-cysteine-containing RING2 domain is encompassed by the *TRIAD supradomain* (IPR044066). In the case of the RBR ligase PARKIN, its activity state is stringently controlled, with both RING1 and RING2 domains being inert in the basal state due to steric occlusion. Activation consists of a complex conformational rearrangement of structural domains (**Fig. 4**) that releases both the E2-binding site and the catalytic cysteine from inhibition^148^. Two of the RBR proteins, RNF31/HOIP and RBCK1/HOIL1, and a third protein, SHARPIN, form the LUBAC complex, which builds linear ubiquitin chains, controlling inflammation, innate immunity, and adaptive immunity^149,150^. HOIL1 has the unusual property of forming oxyesters with the C-terminus of ubiquitin^151^. There is a single RCR ligase in humans, MYCBP2, which as noted above has the unusual property of forming two sequential adducts with ubiquitin prior to transferring ubiquitin to substrate, employing a unique mediator loop for the formation of the first adduct^67^. As with RCR and the RBRs, the RZ ligases RNF213 and ZNFX1 form thioester adducts with ubiquitin. Adducts are formed via the small RZ finger (IPR046439), though this domain does not serve to recruit E2s. For RNF213, the RZ finger is embedded in a so-called E3 shell which, along with and adjacent C-terminal domain, recruits charged E2^68,69^. In contrast, ZNFX1 lacks this structure and uses a variant RING domain to recruit E2^70,71^. The RZ ligases each support innate immunity^68–71^.

### Idiosyncratic E3 enzymes

The E3s described in this survey generally have known or predicted mechanisms for recruiting E2s. We include fifteen proteins as E3s for which the E2 binding component is atypical or unknown (**Table S1**). In vitro autoubiquitination provides good support for E3 function, if the reaction depends on the addition of ubiquitin, E1, and E2. Identification of substrates stabilized by the deletion of a proposed E3 also provides support, particularly if an interaction between the substrate and proposed E3 has been demonstrated. The idiosyncratic E3 enzymes we propose are supported by some or all of this evidence. We include in this group UBR4, which recruits UBE2A/RAD6 via a hemiRING motif which packs into a novel UZI domain (**Fig. 6**)^63^. UBR4 associates with KMCF1^152^, and two recent structural studies have demonstrated that these can combine with calmodulin to form a massive ring structure, recruiting substrates to the interior of the ring^56,57^. In this context, the complex acts as an E4.

### E3s for ubiquitin-like modifiers

The E1 and E2 enzymes for the various ubiquitin-like modifiers (UBLMs) are both functionally and structurally analogous to those of ubiquitin. The UBL ligases show impressive variation, as we have seen in ubiquitin ligases. ISG15, induced by interferon in response to viral infection, was the first UBLM to be recognized as such^153^. The principal ligase for ISGylation in humans is HERC5^154,155^, although several others may contribute^155–157^. SUMO was first discovered as a ubiquitin-like modifier for RANGAP1, targeting it to the nuclear pore complex^158,159^. Myriad physiological functions for SUMO have since been discovered^160–164^ mediated by at least 50 SUMO receptors (**Table S6**)^165,166^. SUMO is ligated to targets via RING variants known as MIZ/SP-RINGs (**Table 1**)^167^, and through non-MIZ ligases (**Table 1**)^168^. NEDD8, while serving critically as a modifier of cullin RING ligases^40^, can modify many other proteins, depending on conditions of the assay^169^. Modification of the CRLs is promoted by the DCN family (IPR005176)^170^, and a variety of bifunctional ubiquitin ligases carry out NEDDylation as well (**Table 1**)^171^.

As described above, ATG8 is conjugated to lipids rather than proteins^27^. This critical activity, which tethers ATG8 to membranes, is mediated by an E3 complex composed of ATG5, which is itself modified with the ULM ATG12, and ATG16L1^172^. The principal target of UFMylation is ribosomal protein RPL26^173^, a modification that promotes the dissociation of ribosomes from the ER translocon, particularly when the translation cycle of secretory proteins is interrupted. This may occur for example as a result of a truncated mRNA, as it will lack a stop codon. This modification is catalyzed by the atypical E3 enzyme UFL1^174^, functioning in complex with UFBP1 and CDK5RAP3^175^. Such translational errors are generally resolved by the RQC pathway, involving multiple ubiquitination events, but the RQC pathway is sterically blocked from acting on ribosomes that are complexed with the translocon on the ER membrane. Thus, UFMylation works closely with the RQC pathway to neutralize translational errors^176–178^.

### Deubiquitinating enzymes

The median half-life of a ubiquitination event in HeLa cells is estimated to be very short–approximately 12 minutes–with some sites being demodified in under one minute^179^. The rapid reversal of ubiquitin modifications is mediated collectively by over 100 dedicated proteases, which are highly differentiated in their specificity, subcellular localization, and regulation. The domains that govern the deubiquitinating enzymes (DUBs) that remove ubiquitin and ubiquitin-like modifiers from their target proteins are large and well-defined, so construction of the DUB Class is straightforward. The USP-type DUBs constitute the largest family of DUBs, and share the InterPro domain IPR028889. This domain also captures PAN2, which functions in mRNA deadenylation rather than deubiquitination. The USP family is elaborated with a variety of other domains, including ubiquitin binding and ubiquitin-like domains, modulating function and substrate specificity. The auxiliary domains that establish Types within the USP Group are listed in **Table S3**. Multiubiquitin chains may be built through any of the seven lysines of ubiquitin or through its N-terminal methionine. On the whole, the USPs do not show strong linkage preferences in their deconjugating activity^180^. The USP17 group comprises a third of the USP-type DUBs, with group members being at least 90% identical. This family was generated by the expansion of tandem repeats on two chromosomes in humans and within the RS447 megasatellite^181^. As with the E2 enzymes, we include catalytically inactive members, of which there are five in the USP Group, three of which are of the USP17 Type. The other flagship group of deubiquitinating enzymes is the much smaller UCH Group, numbering four and covered by the domains IPR057254 and IPR041507. The UCH family has been viewed as dedicated to the removal of small peptides from the C-terminus of ubiquitin, consistent with leaving group restrictions imposed by an active-site crossover loop^182^, but more recently the proteasome-associated UCHL5 has been shown to resolve branched ubiquitin chains^183,184^.

Utilization of the ***MPN domain* (IPR037518)** for deubiquitination traces as far back as archaea such as *C. subterraneum*^185^. In eukaryotes, three multisubunit protein complexes^186^—the lid of the proteasome, the COP9 signalosome, and the translation initiation factor EIF3—contain a pair of JAMM/MPN domain proteins. One is active as a DUB in the proteasome, one as a deNEDDylase in the signalosome, and both as DUBs in EIF3. Of the 15 MPNs we include in the UPS, 10 are active as DUBs, four are included as integral members of active complexes, and one is included as a ubiquitin binding protein. The JAMM/MPNs, uniquely among the DUBs, are metalloproteases rather than cysteine proteases. Most commonly, they act on K63-linked ubiquitin^187^ beautifully exemplified by AMSH^188^, though PSMD14/RPN11, a proteasome subunit, acts at the proximal ubiquitin in a substrate-linked chain^189–191^.

The OTU family shares the *OTU domain* (IPR003323), though OTULIN and its inactive counterpart are identified by *FAM105* (IPR023235). More than half of the OTU family show a strong preference for one or two of the eight ubiquitin-ubiquitin linkages^192^. The MINDY family, a group of four proteins identified by one of two domains (IPR033979, IPR025257), prefer K48-linked ubiquitin, with the exception of MINDY2^187^. ZUP1, in a family of its own, shows strong preference for K63-linked ubiquitin^193^. Uniquely among DUBs, the four members of the Josephin family (IPR012462, IPR049387) show strong preference for ubiquitin esters over ubiquitin isopeptide linkages, and thus provide a largely dedicated pathway for the reversal of non-lysine-directed ubiquitination, notably modifications of threonine and serine^194^.

The remaining DUB families work exclusively or in part on UBLMs. Two of these are structurally related to ZUP1: the UFSP family (IPR012462) and the ATG4 family (IPR046792)^193^. The USFP family functions in deUFMylation^195^, and the ATG4 family is mainly involved in biosynthetic processing of ATG8 family proteins and their delipidation^196^. Most of the SENP family (IPR003653) are deSUMOylating enzymes^197^, but SENP8 acts on NEDDylated proteins^198,199^. The PPPDE pair (IPR008580) are both proposed to be deSUMOylases^200^, though DESI2 additionally functions as a deubiquitinating enzyme^201^. The PPPDE family has also been observed to exhibit cysteine palmitoyl thioesterase activity^202^.

The physiological effects of DUBs have frequently been linked to specific E3s, and in some cases they populate the same complexes (reviewed in ^203^). A DUB can antagonize an E3, or alternatively potentiate an E3 by counteracting suicidal E3 autoubiquitination. A DUB may also fine-tune E3 output by editing ubiquitin chain generated by the E3.

### Ubiquitin and ubiquitin-like binding proteins

As shown in **Fig. 1**, ubiquitination promotes protein turnover through both autophagy and the proteasome, and also promotes signaling events in the absence of any proteolytic targeting. In all cases, it is expected that ubiquitin receptors connect protein ubiquitination with protein fate. We organize ubiquitin binding proteins first by molecular or biological context (described at the Group and Type levels), and then by ubiquitin binding domain (Subtype). We also describe the ubiquitin receptors in an auxiliary table (**Table S6**), ordered first by domain then by molecular context and function.

Close to twenty canonical domains characterize the ubiquitin binding motifs, but even so, only about 60% of the ubiquitin binding elements we list map to InterPro domains. The most frequently used element in ubiquitin recognition is the UBA domain, a three-helix bundle of about 40 amino acids described by the *Ubiquitin-associated domain* (IPR015940) and its related homologous superfamily (IPR009060)^204^. These two InterPro domains, along with an array of niche UBA domains (**Table S6**, Domain Summary tab), identify 92 UBA elements in the human proteome. Given the breadth of this group, members appear in nearly every molecular context and pathway involving ubiquitination. Two contexts are worth special mention: the UBL-UBA proteins that shuttle substrate to the proteasome, and SQSTM1/ p62, a critical receptor for autophagy. Like the UBA domain, the CUE (IPR003892), GAT (IPR004152), and VHS (IPR002014) domains are also composed of helical bundles, the last two are principally involved in biosynthetic subcellular trafficking^205,206^.

UIM elements are simpler than the domains above, consisting of little more than a short stretch of alpha helix. Fewer than half of UIMs that we find are identified by the *Ubiquitin interacting motif* site (IPR003903), the rest being either validated experimentally or identified by sequence and remaining to be confirmed. Though small, UIM domains embedded in proteasome subunit PSMD4/ RPN10 serve as the strongest ubiquitin receptor elements of the proteasome^207,208^. Other simple helical elements involved in ubiquitin recognition include the UBM (IPR025527), UBAN (IPR032419), and MIU elements, the last having no corresponding InterPro domain. Numerous zinc fingers recognize ubiquitin, in aggregate approaching the size of the UBA family. These include *Zinc finger, RanBP2-type* (IPR001876), *Zinc finger, UBP-type* (IPR001607), *Zinc finger, A20-type* (IPR002653), *CALCOCO1/2, zinc finger UBZ1-type* (IPR041641), *FAAP20, zinc finger UBZ2-type* (IPR031490), *DNA polymerase eta, ubiquitin-binding zinc finger* (IPR041298), *Rad18, zinc finger UBZ4-type* (IPR006642), and *NEMO, Zinc finger* (IPR034735). In addition to these, we identify another 12 zinc fingers that are not covered by these domains.

As mentioned above, catalytically inactive members of the E2 family are retained in our survey. Four such variants–UBE2V1, UBE2V2, AKTIP, and UBE2NL– share the canonical E2 domain (IPR000608), and the first two form heterodimers with UBE2N, imposing chain linkage specificity by orienting the acceptor ubiquitin^209,210^, thus supporting DNA repair. The domain IPR000608 falls within a homologous superfamily, *Ubiquitin-conjugating enzyme/RWD-like* (IPR016135), and this encompasses the *RWD domain* (IPR006575) and the *Ubiquitin E2 variant, N-terminal* (IPR008883), both of which we also consider to be signatures for ubiquitin binding. Not only does ubiquitin dock into the active site of E2s, but for about a third of the active E2s, ubiquitin binding on the side opposite the active site, termed backside binding, can promote processivity^211^.

Several structural domains that are not dedicated ubiquitin binders have acquired this property in certain proteins. The *Armadillo-type fold* (IPR016024) encompasses the ubiquitin binding domain found on PSMD2, but this is to date the lone instance of ubiquitin binding for this domain. Other examples include the *CARD domain* (IPR001315), the *SH3 domain* (IPR001452), and the *WD40 repeats* (IPR001680). Atypical ubiquitin binding is not only seen in minority members of domain families, but also among singleton domains and among structures that have not been described in the domain databases (**Table S6**).

In addition to ubiquitin binding elements, our classification also includes three proteins involved in URM association, and more than 50 proteins that associate with SUMO via short hydrophobic elements known as SUMO interaction motifs (SIMs)^165^. SUMO binding is also mediated in a few instances by zinc fingers^212^, WD40 domains^213^, and backside E2 binding^214^. Interestingly, multiple ubiquitin ligases interact with SUMO via SIMs, including RNF4, RNF111, and TOPORS. These are known as SUMO-targeted ubiquitin ligases and ubiquitinate SUMOylated proteins, forming mixed chains and targeting a subset of substrates for proteasomal degradation^215–222^.

### Molecular machines

At the heart of protein turnover by the UPS are two molecular machines, the proteasome^223^ and VCP^224–227^. While there are a number of expansive UPS surveys, few of them include the proteasome, its modulators, and its accessory proteins. We annotate each of these molecular machines and their cofactors as two distinct Classes. Fluctuations in the translation machinery, autophagy components, and the proteasome reflect metabolic states, and all of these should be included in a survey used for evaluating the proteostatic capacity of tissues.

A key underlying feature of protein turnover is how destruction is limited to intended targets. The proteolytic sites of the proteasome are sequestered within an internal chamber of the enzyme, whereas in the autophagy pathway the responsible proteases are sequestered behind the membrane of the lysosome.

The proteasome consists of two major subassemblies, the regulatory particle (RP) and the core particle (CP). The RP, a 19-subunit complex, is also known as the 19S particle, and the CP, a 28-subunit complex, is also known as the 20S particle. Subunits of the CP are described in terms of signature domains (*Proteasome, subunit alpha/ beta*, IPR001353) and assemble into four hetero-heptameric rings, stacked to form a barrel. Gates at the end of the barrel seal the chamber to isolate the proteolytic active sites from the cytosol^228^. These gates can be opened by a variety of activators, the most widely used being the RP. Most importantly, the RP recognizes and prepares substrates for degradation by unfolding them mechanically in the course of injecting them into the CP. The gate-proximal component is a hexameric ring of ATPases (*AAA+ ATPase domain*, IPR003593) with a central pore, which contact substrate and drive unfolding, The ATPase ring is ensconced in a larger structure of non-ATPase subunits which help to situate the ring over the CP gate. As mentioned above, the RP subassembly called the lid is structurally similar to the signalosome and eIF3^186^. Like those assemblies, it contains an intrinsic DUB—PSMD14/Rpn11— which is positioned near the pore of the ATPase ring such that it can trim attached ubiquitin from substrate as it is laced through the ATPase ring and into the chamber of the core particle^191^.

Three additional DUBs dock onto the proteasome, each with distinct locations, substrates, and mechanisms of action^183,184,229,230^. The proteasome also recruits a selection of E3 enzymes to it^231,232^. The activity of the proteasome is modulated by proteasome-specific cofactors that open the gate in the absence of the RP and modulate proteasome activity. Substrate engagement is also promoted by various adaptors that deliver substrate to the proteasome, through either ubiquitin-dependent or independent mechanisms, in the latter case most notably by midnolin^36,233,234^.

Remarkably, more than 25 kinases, phosphatases, and other enzymes modify the proteasome and promote or suppress its activity^235^. We do not include them as UPS components, as these modifying enzymes are not strictly required for proteasome activity, nor are the enzymes dedicated to the proteasome. Nonetheless, the physiological impact of these modifications can be significant, even if the dynamic range of the activity changes is in the realm of fine-tuning.

Although the proteasome is equipped with the capacity to unfold and translocate proteins into its proteolytic chamber, it cannot extract proteins from intracellular membranes or, typically, protein complexes. VCP fills this void, for example extracting proteins from and through membranes, thus preparing them for degradation by the proteasome. In this way, VCP promotes ERAD and degradation of outer mitochondrial matrix proteins. Soluble proteins can also be poor proteasome substrates due to an inability of the proteasome to mediate their unfolding, and in such cases VCP may carry out the initial unfolding step.

Like the proteasomal ATPases, VCP is a hexameric ring of ATPases formed around a central pore. Each VCP subunit has two iterations of the AAA+ domain, giving rise to a double ring structure. N-terminal domains emanate from each monomer, and serve to recruit adaptor proteins such as UBX domain proteins, a specialized type of UBL-domain protein. Several of these adaptors target ubiquitinated substrates. VCP lacks an intrinsic DUB, but recruits a number of DUBs, as well as a number of E3s. Substrates are unfolded by being passed through VCP’s central pore before the proteasome can act on them. In addition to assisting the proteasome, VCP participates in myriad cell processes, including extracting proteins from chromatin^236^, promoting ribosome quality control^237^, reassembling the Golgi following mitosis^238,239^, and promoting the clearance of ruptured lysosomes by autophagy^240^.

### Comparison of Compendia

The central motivation of our work was to create a reference resource for further studies of the UPS. We therefore designed the survey to be comprehensive, to embed community feedback in its delineation, and to provide a hierarchical structure for organizing UPS genes. Parts of the UPS have already been the focus of prior annotation efforts by other groups. Comparison to those efforts allows us to assess the robustness of our annotations through convergence, as well as to identify conceptual boundaries that emerge from differences in classification.

Our survey is the first to include all eight Classes of the UPS, and it continues a decades-long effort by many in the field to delimit this vast biochemical system. Such efforts, including ours, have relied to variable extents on assessment of the literature, text mining, domain searches, and the elaboration of earlier annotations (**Table S7**, Collections tab). We selected a subset of these for comparison with the current study^114,241–248^.

The scopes of prior studies and their relationships to each other are shown in **Fig. 9A**. We compared the aggregate membership of these nine UPS sets to the UPS proteins of the current study (**Fig. 9B**, Venn diagram I). In this comparison, there were 167 members unique to our set, roughly half being ubiquitin binding proteins. Surprisingly, 819 proteins previously proposed as UPS components were not included in our collection. The aggregate set was refined by removing proteins that do not carry specific annotations or were computationally derived (see Methods) and again compared to the UPS members in the current study (**Fig. 9B**, Venn diagram II). This relatively simple operation accounts for 548 of our exclusions, but this leaves 265 additional exclusions, reached typically through more ad hoc considerations, and 235 of these had been proposed to be ubiquitin ligases.

**Figure 9.**
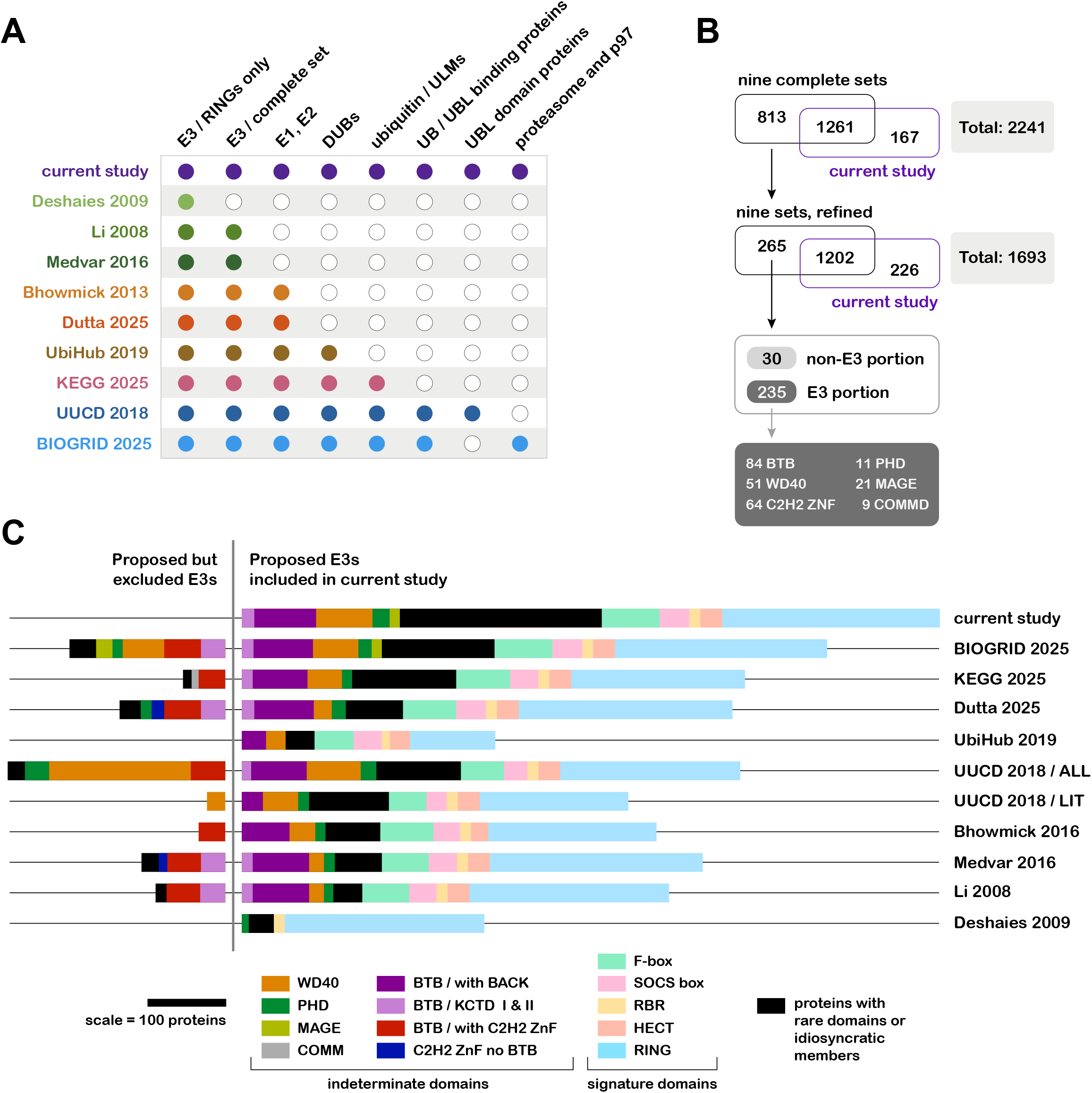
Comparison of UPS annotations including E3 ligases. (**A**) The scope of nine previous UPS compilations (**Table S7**) chosen for comparison to the current study is shown. Colored dots represent inclusion of the indicated category in related survey. (**B**) An aggregate of the nine earlier studies compared with members of the current study (first Venn diagram). The aggregate set was refined, removing proteins that do not carry specific annotations or were computationally derived (see Methods), and again compared with the current study (second Venn diagram). The excluded proteins in the second analysis were partitioned into those proposed to be E3 ligases and not (third diagram), and evaluated for enriched InterPro domains (fourth diagram). (**C**) For each compilation, the number of proteins excluded or included relative to the E3 set in the current study is shown by scaled bars. InterPro domains which are not sufficient for inclusion as E3s (indeterminate domains) are shown in more saturated colors, and InterPro domains that are sufficient for inclusion (signature domains) are shown in pastels. Minor domain types (<9) were omitted from the graph.

We conducted a domain analysis on the set of proposed E3s that we excluded from our collection, and found that more than 80% shared at least one of six InterPro domains: MAGE, COMMD, PHD, BTB, C2H2 zinc fingers, and WD40 (**Fig. 9B**, bottom). While we do include MAGE RING cofactors in our survey, not all MAGEs are thought to have this function^249^. All proteins with COMMD domains are included in KEGG, but we find it more likely that only COMMD1 is involved in E3 regulation^250^. A handful of the 88 PHD domain proteins have been identified as variant RING ligases, but participation of PHD domains (IPR001965) in E2 binding is thought to be the exception rather than the rule^114^. A group of 101 excluded E3s contain BTB domains (IPR000210), C2H2-type zinc fingers (IPR013087), or both. BTB domains dock into the adaptor-binding region of CUL3, but this domain alone is not necessarily sufficient for docking. In BioPlex 3.0, CUL3 recovered BTB-BACK and KCTD-type I BTB proteins (a third of each group), but was not found with any of the 27 KCTD-type 2 BTB proteins nor with any of the 49 BTB proteins containing zinc fingers. The entire repertoire of BTB proteins was included in an initial catalog (**Fig. 9B**)^241^, but this now seems overbroad. Moreover, zinc finger clusters found in BTB proteins are structurally distinct from zinc-binding RING domains, but given that they are appended to BTB domains, they may have come to be construed to be E2 recruiting domains over time.

The last group of excluded E3s is the WD40 set, numbering 51 members. The WD40 domain is found among the receptors for most most types of cullin RING ligases, and in the context of F-box proteins, SOCS box proteins, and BTB domain proteins, the WD40 element does not contact the cullin adaptor. For CUL4A/CUL4B substrate receptors, the WD40 domain engages adaptor DDB1 but does not constitute the entire interface. The putative CUL4A/ CUL4B receptors were identified in a parallel set of proteomic experiments 20 years ago. Some hits turned up in only one of four screens^93,94^, were not subsequently identified by BioPlex, and do not model well with DDB1. Of these, 14 have been included in previous annotations, but we exclude these give their lack of validation. Even so, we consider these to be borderline cases at present, with reasonable arguments both for and against inclusion. Overall, the inclusion of E3s that we exclude may be rationalized to a large extent as the consequence of overbroad application of domains as sufficient for inclusion as E3s.

To visualize the relationships between the E3 component of our annotation and earlier E3 sets, we applied our domain analysis graphically, and to both the excluded and included portions of the sets (**Fig. 9C**). Over time, the scope of E3s identified by canonical domains has expanded (**Fig. 9C**), which is consistent with the continuing elaboration of domain assignments in InterPro. The BioGRID E3 collection most closely resembles ours, with 718 shared components and 144 E3s that are unique to our set. Roughly 80% of these differences correspond to E3 components that are idiosyncratic or are included by virtue of rare domains (**Fig. 9B**, black bars), with the other 20% corresponding to a broader application of canonical domains. Among the more prominent annotations unique to the present survey are PRAME CUL2 receptors, and noncatalytic subunits of idiosyncratic ubiquitin ligase complexes. In **Table S7**, we compare our annotation to previous annotations on a protein-by-protein basis. Pairwise comparisons between the current study and earlier studies are also made, and can be found in **Table S7**.

Two studies were published contemporaneously with submission of the initial version of this manuscript: Szulc & Pokrzywa (2026)^251^ and Chua et al. (2026)^252^. The former focuses on the cullin RING ligase substrate receptors, the latter on the whole E3 class. We compare the applicable sections of our survey to these catalogs in **Fig. S3** and in **Table S8**. In comparison to the E3-ome of Chua et al. (2026)^252^, the E3 class in our survey is broader by 206 components (**Fig. S3A**). Half of these derive from our more expansive sense of what constitutes an E3. We include all subunits of E3 complexes, rather than a subset, as functional forms of E3s typically require all subunits to be present. We also include all recognized substrate adaptors—components which associate with fully-formed E3 ligases and augment or alter their substrate specificity or affinity. The other half of proteins that are unique to our study fall into categories represented in both studies: RING and variant RING ligases, idiosyncratic ligases, UBLM ligases, and substrate receptors for cullin RING ligases. The 15 E3-ome components that we exclude from our study are predominantly proposed CRL substrate receptors (**Fig. S3B**). Likewise, Szulc & Pokrzywa (2026)^251^, which catalogues 267 CRL substrate receptors, proposed 21 CRL substrate receptors not present in either the current study or in Chua et al. (2026)^252^.

No two studies of the scope of the UPS, or of the diversity of ubiquitin ligases, have been fully in agreement. We have summarized the differences between the present and numerous others in **Fig. 9, Table S7, Fig. S3**, and **Table S8**. Given the multiplicity of previous studies, it is not feasible to fully account for the different conclusions reached here, for lack for space. Briefly, while the known breadth of the UPS is continually expanded through new experimental work, differences from earlier studies arise more from the marked improvement of predictive approaches in the last few years, with the development of AlphaFold and related tools such as the Predictome. Ongoing proteome-wide interactome studies are also broadening our perspective on the UPS, but can in general only be interpreted in light of additional data. For example, novel E3 components cannot be assigned from interactome studies, since they do not resolve ligase components from ligase substrates. Also, some protein family-based classifications have proven to be overly reductionist. Once multiple BTB proteins were found to function as CUL3 receptors, it was often assumed to be an E3 signature and a sufficient basis for classification. Subsequent experimental resolution of subtle requirements for BTB docking onto CUL3 created a framework for defining true E3 components from the broader BTB family.

We welcome community feedback on the composition of the Proteostasis Network, submitted to. The exact composition of the PN will be revised as new data become available, and updates will be posted on the Proteostasis Consortium website (https://proteostasisconsortium.com). An interactive web resource is now available for exploring the contents of the Human Proteostasis Network (https://proteostasisnetwork.streamlit.app/), which allows both user-directed exploration and data downloads.

## DISCUSSION

To date, no two surveys of the UPS have arrived at very similar censuses of its components. This reflects the exceptional breadth and complexity of the system, as well as the impact of significant ongoing advances in the field. Even more so perhaps, the methods used to define UPS components have led to different outcomes. At the same time, some surveys of the UPS have drawn from earlier surveys and in doing so have propagated what we consider to be assignment errors. We have avoided metrics such as frequency of annotation to favor inclusion, as this will result in the amplification of early misassignments. Although in many surveys the noncatalytic subunits of E3 enzymes are omitted, they are critical components, and we therefore include them in this survey, placing them in their relevant context. We also include cofactors of E3s that, while not intrinsic subunits, are required for the modification of particular substrates.

We made extensive use of recently developed tools and databases to aid in defining the UPS: the BioPlex 3.0 database^253^, AlphaFold3^254^, and the Predictomes project^73^. While using these tools, no assignments were made using machine learning per se.

We emphasized a domain-based approach for inclusion in which the functional features of the domains help to undergird and explain why a protein should be considered a component of the UPS. We also note these domains explicitly, and comment on domains that appear frequently among UPS proteins but do not predict function narrowly. Especially for candidate members lacking obvious domains to associate them within the UPS, we scrutinized the literature before placing a proposed UPS component in our survey via entity-based criteria. In using primarily a domain-based approach for classification, we have included proteins that have not been studied in any detail biochemically–further experimental work on these little-known proteins should be a fertile area for new insights into the UPS and solidify our understanding of their place in the UPS.

In addition to providing a comprehensive survey of UPS components, but we also arrange these in a five-tiered taxonomy that allows for a clear understanding of the position of each protein in the pathway as well as highlighting proteins with similar activities and structural features. This survey is also unusual in covering every aspect of the UPS, including ubiquitin binding proteins, proteasomal proteins, and proteasome-associated cofactors. In covering all aspects of the UPS systematically, we encountered a complexity incompatible with single annotations: for example, UBL domains and ubiquitin binding domains each occur in numerous enzymes, including E2s, E3s, and DUBs. Likewise, while some proteins that associate with molecular machines are dedicated to their maturation and modulation, others also have functions in the UPS beyond their collaboration with molecular machines. These cross-annotations are noted in the **Table S1**, and described in **Fig. 10**.

**Figure 10.**
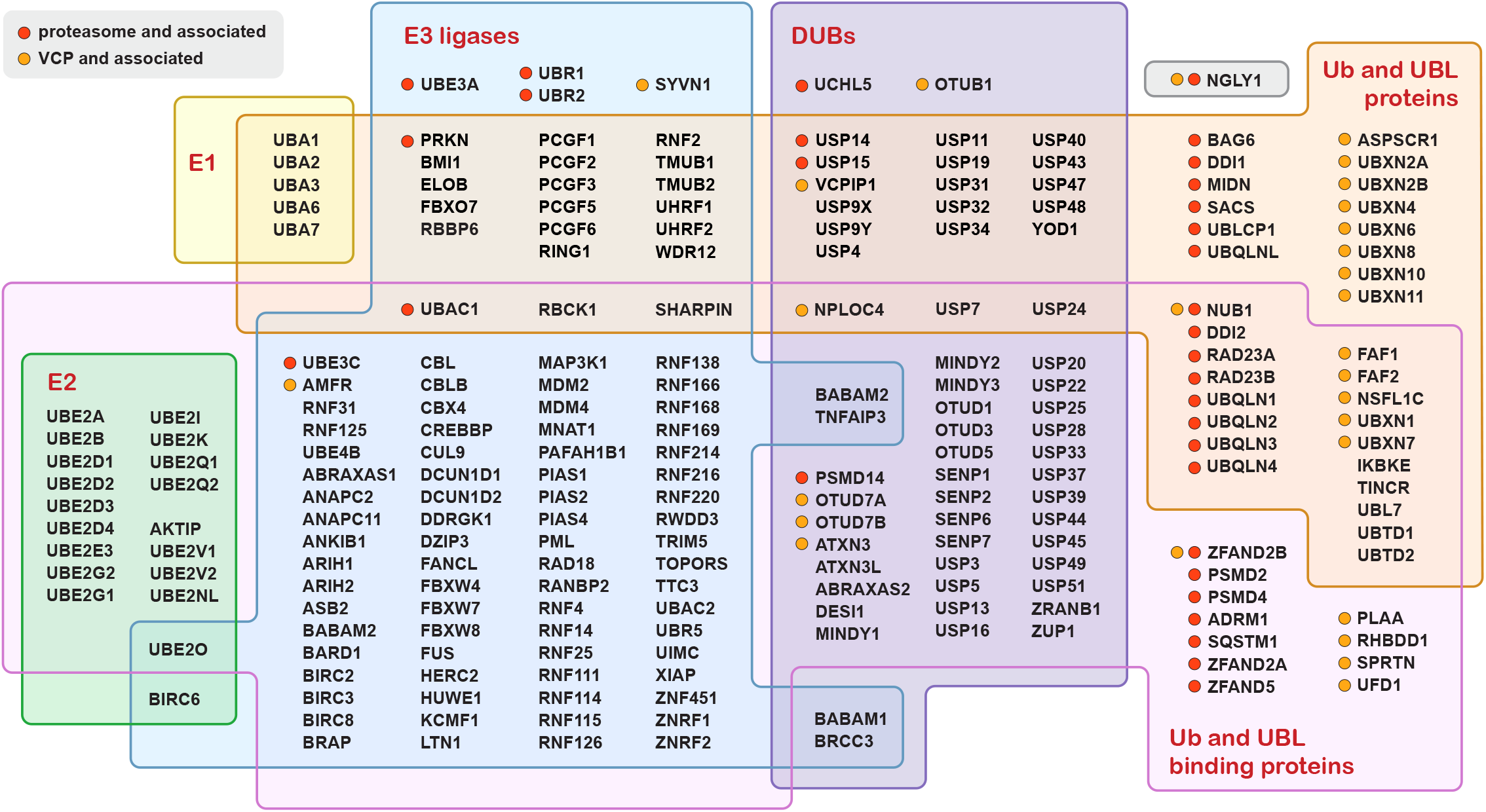
UPS genes with annotations in more than one Class within the UPS Branch. Venn diagram shows six of eight UPS classes: E1 activating enzymes (yellow), E2 conjugating enzymes (green), E3 ligases (blue), deubiquitinating enzymes (purple), ubiquitin and UBL proteins (orange), and ubiquitin and UBL binding proteins (pink). Proteins associated with the molecular machines are indicated with red and orange dots. Only the UPS proteins annotated in more than one Class, comprising 15% of the survey, are shown.

The present UPS survey was developed in concert with surveys characterizing other parts of the PN: translation, protein folding, organelle-specific proteostasis, and autophagy^11,12^. A remarkable 200 UPS genes, approximately, are cross-annotated in other Branches of the PN (**Fig. 11**). These cross-annotations arise from known features of the pathways involved, most notably the shared ubiquitin-dependence of much proteasomal and autophagic degradation. Consistent annotation of the whole of the PN using a uniform taxonomy should provide a powerful groundwork for the field of proteostasis and its fundamental roles in human health. Among the many applications for this survey, the most straightforward is to improve the precision of enrichment analyses performed on global datasets at the genomic, transcriptomic and proteomic levels. To understand the overall behavior and functionality of the UPS in a given differentiated tissue, age, metabolic state, or stress condition, a reference assembly of system components that is as accurate and complete as possible is indispensable. Given that this study provides by far the most extensive UPS survey on record, it will be interesting both to filter new data through this analytic framework, and to use it to reassess previously published genomic, transcriptomic, and proteomic datasets for previously undetected or incomplete UPS signatures, including poorly understood disease-associated genetic loci, hits from large-scale CRISPR screens, and targets screens involving potentially therapeutic small molecules.

**Figure 11.**
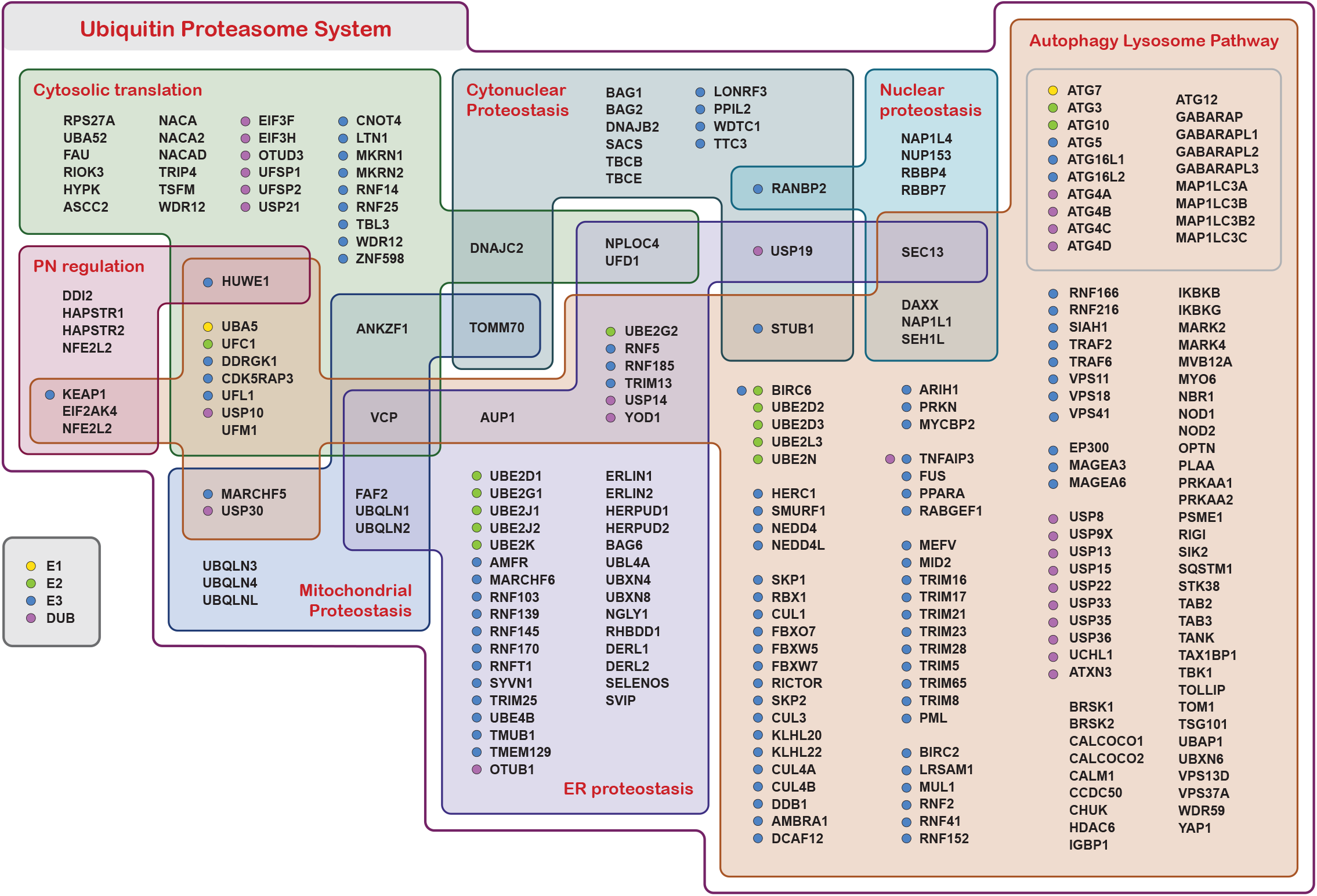
UPS genes annotated in other Proteostasis Network Branches. Proteins with annotations in the Ubiquitin-Proteasome System and in at least one other Proteostasis Network Branch are shown, listed with their official gene symbols. Colored dots indicate functions in the UPS pathway as noted in the key at lower left (grey box). Components without dots fall within the other four UPS Classes: ubiquitin and UBL proteins, ubiquitin and UBL binding proteins, proteasome and associated proteins, and VCP and associated proteins). The genes corresponding to the ubiquitin-like modifiers Atg12 and Atg8 (with its homologs) in a grey box within the ALP sector at upper right, along with the enzymes that mediate their conjugation and deconjugation.

It will be important to survey the UPS and more generally the PN in model organisms as well, and this is in progress. Previous work has shown that, remarkably, the UPS was present in a relatively mature form as early in evolution as the Last Eukaryotic Common Ancestor (LECA): It has been estimated that approximately 95% of UPS gene families were represented in the LECA^255^. Therefore, the UPS clearly took shape and assumed great complexity during eukaryogenesis, that is, prior to the LECA. During this unique evolutionary period, the proteome of the cell must have been in tremendous flux, resulting in stringent new demands on both protein and organellar quality control. Adaptation to these conditions during eukaryogenesis may have driven the elaboration and progressive refinement of the UPS and of the proteostasis network as a whole.

## METHODS

### Counting UPS Genes and Proteins

For the UPS, there are five proteins encoded by more than one gene: Q0WX57 is encoded by 7 genes (*USP17L24, USP17L25, USP17L26, USP17L27, USP17L28, USP17L29, USP17L30*), C9J1S8 is encoded by 2 genes (*TRIM49D1* and *TRIM49D2*), P43356 is encoded by 2 genes (*MAGEA2* and *MAGEA2B*), Q16637 is encoded by 2 genes (*SMN1* and *SMN2*), Q9GZY0 is encoded by 2 genes (*NXF2* and *NXF2B*). Therefore, the UPS gene count is 10 greater than the UPS protein count, which is 1402. When comparing our survey to previous annotations, we compared Uniprot IDs rather than Gene IDs, so numbers are expressed in terms of proteins rather than genes.

### Counting E3 enzymes

To obtain a count of the E3s, we considered all unique E3 complexes. For the cullin RING ligases, each substrate receptor (n=315) and each substrate adaptor (n=11) define a unique E3. CUL4 adaptor AHR-ARNT-TBL3 forms a complex, and is considered to be a single adaptor. The CUL4 scaffold DDA1 is not counted because it is found in the context of substrate receptors already counted. Most cullins, the RBX proteins, the cullin adaptors, and the cullin regulators are all subsumed in this number. We count CUL7 since it appears to have the capacity to function not only with substrate receptors but also autonomously. CUL9 is likewise counted (n=2).

For the RING ligases, we count the TRIMs (n=74) but not the ringless TRIMs, nor TRIM heterodimers, since it is uncertain how many heterodimers actually form, and so TRIMs may be undercounted. For the remaining RINGs, we count all individual RINGs (n=230). We also count the RING adaptors individually (n=19), as recruitment of each adaptor function defines a new complex. We do not consider that an adaptor may associate with more than one RING, possibly undercounting. For the idiosyncratic RING complexes, each complex of proteins is counted once (n=11), with the exceptions of the APC, which has two known adaptors (n=2) and CTLH, which has six known adaptors (n=6). The CTLH adaptors dock at multiple sites and may therefore work in numerous combinations, and we do not consider these, which is another likely source of undercounting. The RING variants are counted individually: UBOX (n=8), PHD (n=7), and C4C4 (n=1).

For the smaller classes of E3s, we count the RBRs individually (n=14), and also count LUBAC (n=1), as the two RBR proteins it includes may have individual functions. The RCR (n=1) and the RZ ligases (n=2) are considered individually. The HECT ligases are each counted individually (n=28) and their adaptors are counted additionally (n=13), and each of these is only counted once though adaptors may bind multiple HECTs, representing another undercount. The HECT activators are not counted. The E2-E3 fusions are counted individually (n=2) as well as the UBE2O adaptors (n=2). There are three idiosyncratic E3 enzymes that are counted individually (n=15) and one idiosyncratic complex (n=1). We additionally count the E3s for UBLMs and their adaptors (n=27). The total for the minimum number of E3 enzymes in human is 792.

### Identification of UPS proteins

To generate a comprehensive accounting of the UPS (**Table S1**), we first assessed well-characterized UPS protein families for the presence of shared InterPro domains, which we term signature domains. We also screened the InterPro domain database of more than 50,000 entries for additional signature domains. By querying UniProt with these signature domains (**Table S3**), we identified additional family members. In addition to this domain-based approach, we consulted the literature at length, evaluating review articles and the primary literature with a focus on new UPS developments (**Table S1**). A simple keyword query of UniProt was found to be far less effective than probing with our curated InterPro domains; searching with “ubiquitin OR proteasome” retrieved more than three thousand proteins, including myriad proteins not ultimately included in our survey and missing hundreds that we ultimately included. After the core of our survey was assembled, we consulted a range of existing databases, including BioGRID, KEGG, and Gene Ontology (focusing on the Molecular Function hierarchy). We also evaluated earlier UPS annotations, Bhowmick et al. (2013)^243^, Medvar et al. (2016)^242^ and Zhou et al. (2018) (UUCD 2.0)^244^. When available, we consulted citations relating to specific genes within these resources. To evaluate E3s lacking signature domains, we weighed most heavily evidence showing ubiquitination in a purified in vitro system dependent on the proposed E3, and these references are included in **Table S1** (Main tab).

### Assignment of Signature and Auxiliary Domains

Our survey of more than 1430 proteins has been parsed in terms of protein function, and we link function as often as possible to specific protein domains. InterPro and CDD domains for member proteins were used to develop the taxonomic structure of our survey. To ensure a unified description, the catalog was evaluated near the end of its construction with the profiles present in UniProt 2025_04, (released on October 14, 2025). Values for InterPro domains, PDB structures, and protein existence correspond to those present in UniProt 2026_01 (released on January 27, 2026). The InterPro IDs were matched to the entries in InterPro 107.0 (released October 15, 2025), with the last entry being IPR059259. In a handful of cases, InterPro domains that had not yet been incorporated into UniProt were identified and included in our annotation.

The Class level is generally too broad to be described by a single unifying InterPro domain (with some exceptions, such as E1 enzymes). Most proteins are annotated with “signature domains” which place them in a particular Class-Group or Class-Group-Type classification. We use “auxiliary domains” to further refine signature domain groupings. We also implement auxiliary domains to note features that appear in multiple Group-Type listings. For VCP-associated proteins and ubiquitin binding proteins, we implement signature domains differently. Rather than using them to designate broader categories, we use signature domains to indicate the functional elements that confer VCP or ubiquitin binding, which we list at the Subtype level.

When InterPro domains are used to represent function, there will occasionally be wholly overlapping domains relating to the same functional element. For a given set of proteins, we chose domains which encompassed the largest number, and then the next largest until we had a complete set of domain designations. We gave preference to InterPro Domains and Repeats over Homologous Superfamilies and Families. Domains that were overbroad for our purposes were avoided where possible. Where unavoidable, we show these domains in parentheses. The specificity of each domain is described in **Table S3** by listing the number of appearances in the UPS, the PN, and the human proteome.

Ideally, all entries sharing the same Class-Group or Class-Group-Type annotation would also share the same signature domain, but there are deviations from this ideal in our survey. For example, the F-box proteins nearly all share InterPro domain IPR001810, but some are only identified with the cognate homologous superfamily IPR036047. In some cases, related homologous superfamilies are overbroad, but not for the F-box proteins. We consider both IPR001810 and IPR036047 to be definitional for inclusion as F-box type CUL1 receptors. In both cases, the signature domains relate to a structural element that docks into the CUL1-SKP1 complex. Some annotation groups include members related by function but not structure. RICTOR shows good evidence of being a CUL1 substrate receptor^256^, but is not an F-box protein. We include it in the survey, but we do not assign it a signature domain.

F-box proteins are additionally annotated with auxiliary domains, including LRR repeats (IPR032675), *WD40 repeats* (IPR001680), and *FBA domains* (IPR007397). We also give auxiliary domains to singleton Subtypes when that domain appears in other parts of the annotation. Only one F-box protein contains a SPRY domain (IPR001870), but there are more than 70 proteins in the UPS that contain this domain. Occurrences of auxiliary domains are not always noted. Some of the ATGylation E3s contain WD40 repeats, but the grouping is small enough that we do not apply an auxiliary domain to differentiate its members.

In some cases, auxiliary domains are implemented without signature domains. While most of the CUL4 substrate receptors have WD40 domains, a signature domain for CUL4 substrate receptors should ideally predict the interaction with DDB1, but no InterPro domain captures this feature. Still, the enrichment of WD40 proteins among CUL4 substrate receptors is a significant feature of the grouping (appearing at a frequency of 44-fold higher than in the proteome as a whole) and is described with an auxiliary domain. Idiosyncratic E3 enzyme complexes are frequently described without signature domains for two reasons: first, InterPro domains that predict complex formation for these complexes are rare, and second, characterization with a domain that occurs only once in the proteome is of little use in our taxonomy.

InterPro domain designations are continually expanded, with thousands of entries being added each year. We anticipate that new InterPro domains will be assigned for known functional motifs that are characterized in the literature and in this survey.

### Structural Analysis and Modeling

Determinants of cullins involved in docking adaptors and substrate receptors were evaluated by analyzing more than 20 solved structures involving all of the canonical cullins (**Table S4**), though CUL4B was used to represent CUL4A and CUL4B. Structures were evaluated using PyMOL. Residues at the cullin-adaptor and cullin-receptor interfaces were identified by screening for residues in these proteins falling within 4Å of their binding partners. The relative position of binding determinants on helices 1-4 were determined and mapped onto a cartoon of the four-helix bundle (**Fig. 7B**).

To establish support for inclusion of the PRAME family as CUL2 receptors, we used AlphaFold3 to model representative PRAMEs with the core components of CRL2. Modeling the CRL2 core complex alone, which contains CUL2 and its heterodimeric adaptor ElonginBC, generated an ipTM score of 0.81. To establish an ipTM range for authentic CRL2 receptors, we modeled all substrate receptors for which structures had been obtained experimentally (VHL [PDB:4WQO], APPBP2 [PDB:8JAL], FEM1B [PDB:8JE1], FEM1C [PDB:8PQL], and LRR1 [PDB:7PLO], resulting in ipTMs ranging from 0.63 to 0.77. PRAME family representatives PRAME, PRAMEF4, and PRAMEF10 modeled with ipTM scores of 0.66, 0.68, and 0.64, respectively, solidly within the range seen for established CRL2 receptors. In these models, the PRAMEs docked at the same surfaces of CUL2-ElonginBC as the established CUL2 receptors, which provides additional support for the PRAMEs as true CRL2 substrate receptors. The PRAME family is strongly conserved in the CUL2/ElonginBC binding element, which spans the first 120 residues.

To evaluate the BTB-BACK proteins for CUL3 association, we consulted the models present in Predictome^73^. Where BTB-BACK-CUL3 models were not already available, we generated models using AlphaFold3 using full length proteins and the default parameters. We construed BTB-BACK proteins as modeling poorly with CUL3 if the modeled pair had ipTM values lower than 0.6, which AlphaFold3 sets as the lower limit for potentially correct models. In most cases, BTB-BACK proteins that modeled poorly with CUL3 docked at sites other than the N-terminal four-helix bundle.

To evaluate the proposed CUL4 receptors identified through affinity screens^90,91,93,94^, each one was modeled with DDB1 using AlphaFold3, the default parameters, and full length proteins (for TOR1AIP2, the 131-residue splice variant was modeled). Proteins indicated to be CUL4 substrate receptors by biochemical evidence were likewise modeled. The resulting ipTM scores showed a bimodal distribution, with very few falling between 0.25 and 0.75. The results are presented in **Fig. S1**.

### Comparison of current study with earlier annotations

#### Data set inclusion and preparation

In order to compare the current survey with earlier studies, we distilled our annotation such that all features of each protein were represented in a single table row. Annotations for the E1, E2, E3, DUB, proteasome, and VCP Classes were truncated, then combined into a thumbnail annotation (**Table S7** [Index tab]), and the number of initial annotations was noted. UBL domains and ubiquitin binding domains are noted in separate columns, and also represented in the thumbnail annotations for proteins having only these features. Some resources are static and contain single annotations for each entry, including Li et al. (2008)^241^, Deshaies and Joaziero (2009)^114^, Bhowmick et al. (2013)^243^, Liu et al. (2019) (UbiHub)^245^, and Dutta et al. (2025) (Human E3 Ligome)^246^. Medvar et al. (2016)^242^ is similar to the aforementioned, though E3s are also annotated as definitive or related, and that information is retained in the relevant tab. Zhou et al. 2018 (UUCD 2.0)^244^ includes multiple annotations for some entries, and the scope includes both UBL domain proteins and ubiquitin binding proteins. We compressed that annotation, applying the approach that we used for the current study. The UUCD 2.0 resource distinguishes between annotations that are derived from the literature and those that were arrived at computationally, and this notation is retained. BioGRID is an actively updated resource, and UPS components are profiled as part of a themed curation project (see **Table S7**, [Collections tab] for link). The human component of the collection was downloaded on December 26, 2025. Likewise, the KEGG resource is actively updated, and the UPS is covered by KEGGBRITE ko04121 (see **Table S7** [Collections tab] for link). That collection was downloaded on January 5, 2026, and the human genes were identified and carried forward for analysis. KEGG uses a four-tiered taxonomy for annotation, and compressed annotations were prepared as above. The original four tiers are included in the relevant tab in **Table S7** (though merged for the small number of genes with multiple annotations). Lastly, UniProt was queried with the search string “E3 ubiquitin-protein ligase” on December 26, 2025.

#### Identification and Classification of Excluded Proteins

We compared the proteins annotated in the current study with an aggregate set comprised of the nine annotation efforts listed above (**Fig. 9A**) combined with the results of the UniProt keyword search. There were 1273 proteins found in both sets, 154 proteins found only in the current study, and 928 proteins excluded from the current study. We divided these excluded proteins into four tiers, with Tier 1 encompassing the most frequently annotated genes and Tier 4 the least. Tier 4 proteins (115) had been defined as such solely on the basis of the UniProt keyword search results. Tier 3 proteins (298) in the aggregate set correspond to the “unclassified” and “no annotation” categories from Dutta et al 2025^246^, and are not included elsewhere. Tier 2 proteins (250) correspond to proteins in the computational portion of the UUCD 2.0 set, and not present elsewhere (except for those discussed above in Dutta et al [2025]^246^). Tier 1 proteins (265) comprise the excluded portion from the narrowest comparison (**Fig. 9B**, second Venn diagram). In summary, borderline cases for exclusion are most likely found in Tier 1, whereas stronger arguments for exclusion apply to Tiers 2-4. All 928 of the excluded proteins are listed with tier designations in **Table S7** (Excluded tab). Annotations from the earlier studies are also included for comparison. Note that in **Fig. 9B**, the Venn diagram I reflects the status of the comparison after the exclusion of the Tier 4 proteins. Venn diagram II reflects the status of the comparison after the exclusion of the Tier 3 and Tier 2 proteins.

The process of narrowing the aggregate set revealed lightly annotated proteins that we include in our survey. Group A includes 167 proteins that were not annotated in any of the earlier studies we compared. Group B includes 24 proteins that were described in Dutta et al. (2025)^246^ as “unclassified” or “no annotation” and do not appear in earlier sets. Group C includes 35 proteins that are included in the UUCD 2.0^244^ computational set but do not carry other annotations beyond Dutta et al. (2025)^246^. The remaining 1212 proteins are included in Group M, each with at least one and typically multiple annotations. The members of each group and their relationship to the current and previous annotations are noted in **Table S7** (Included tab). Borderline cases for inclusion are most likely to be found in Group A, stronger cases for Groups B and C.

#### Pairwise and Aggregate Comparisons

Each of the resources we chose for analysis is characterized in a unique tab in **Table S7**. Previously published catalogs are presented using the categories used by the authors of each previous study, and the annotations themselves hew as closely to the original while striving for brevity. UniProt IDs and official gene symbols have been updated as appropriate. The Index tab provides the master list for the analysis, and encompasses both proteins included in and excluded from the present survey. The Included and Excluded tabs relate information in the Index tab to earlier studies. The Excluded tab presents all UPS proteins proposed in earlier studies but excluded from our set. In this tab, the 387 proposed E3s are presented first. Of these, 331 contain domains showcased in **Fig. 9B** and are sorted by these domains. A group of 12 proposed UBL domain proteins and 103 ubiquitin domain proteins follows the E3 section. Following this is a group of 16 proteins present in BioGRID that do not fall into the above categories. Most of these are ERAD proteins, and they are all included in the ER Proteostasis Branch of the network presented in one of the related Proteostasis surveys^11^. Most ERAD pathway components, such as ubiquitin ligases, are included in the present survey, though some, such as upstream components of the ER lumen, are only given a in the survey of the ER Proteostasis Branch^11^. The remaining 407 proteins are listed last, with a subset of 146 proteins sorted by shared domains.

#### Comparison of the current study with the most recent annotations

Following the initial submission of this manuscript, a new catalog of ubiquitin ligases was published^252^ as well as a more specific catalog of substrate receptors for cullin RING ligases^251^. Ubiquitin ligase components from the current study were compared to the E3-ome^252^, with 656 shared components, 206 specific to the current study, and 15 specific to the E3-ome catalog (**Fig. S3A**). An aggregate set of cullin RING ligase components from the catalogs noted above was compared to that of the current study, with 295 shared components, 52 specific to the current study, and 36 excluded from the current study (**Fig. S3B**). The two recent catalogs are each compared to the current survey on a gene-by-gene basis in **Table S8**. Ubiquitin ligase components present in the current study but excluded from the E3-ome are listed in **Table S8** (divergence_1 tab). Cullin RING ligase components proposed in the recent catalogs but excluded from the present study are listed in **Table S8** (divergence_2 tab), and annotated with information given in the Index tab.

## Supporting information

Table S1. Survey of the Human Proteostasis Network 4.3.3

Table S2. Overview of the UPS Survey

Table S3. InterPro Domains of the UPS

Table S4. Protein Structures

Table S5. Predictions of E2-E3 pairings

Table S6. Ubiquitin and UBL Binding Proteins

Table S7. Comparison to Previous UPS Catalogs

Table S8. Comparison to 2026 Ubiquitin Ligase Catalogs

Figure S1. DDB1 and CUL4A/4B Associated Proteins Modeled with DDB1

Figure S2. A Subset of BTB Proteins Expected to Function as CUL3 Partners

Figure S3. Comparison of the Current Study to 2026 Ubiquitin Ligase Catalogs

## GUIDE TO SUPPLEMENTARY TABLES

### Table S1. Human Proteostasis Network Annotation

The complete annotation of the Human Proteostasis Network is presented in the Main tab. The annotation of the UPS Branch begins at Row 2874 in the Main tab, and is additionally presented in the UPS_only tab. Detailed information about the tabs and fields is given in the README tab.

The five-tiered taxonomy presented in this catalog supports fine-grained enrichment analyses for transcriptomic and proteomic work. The UNIQUE tab offers a repository of indexing values that can be used to relate external data sets, with Gene IDs and UniProt IDs being present in both the MAIN and UNIQUE tabs. The MAIN tab includes columns detailing annotation numbers and associated branches, while the ANN_counts tab collects components with similar profiles in sets. Ordering the MAIN tab by Gene Symbol allows a quick visualization of all annotations for a given gene, and the original order can be recaptured easily with the ORDER tab. The TALLIES tab shows the scope of each Branch and Class.

### Table S2. Overview of the UPS Survey

Classes, Groups, Types, and Subtypes are displayed with gene counts for taxonomic levels. This table is an elaboration of **Fig. 3**, but is more concise than the complete annotation (**Table S1**). This table is not meant to be used computationally, but instead serves as a useful summary of the UPS annotation. For most users, this table will be an optimal entry point. The Class, Group, and Type designations from Table S1 are employed, and the presentation order from Table S1 is maintained. An index is provided on the first of six pages, and coherent sets are grouped by page or column.

### Table S3. InterPro domains of the UPS

Signature domains, distributed domains, and auxiliary domains of the UPS are listed. Domains are described in terms of the Classes or Groups they annotate. Frequencies of occurrence in the UPS, the PN, and the proteome are given for each domain.

For the UPS in particular, inclusion of proteins in the survey relies heavily on signature domains, and those domains are present in MAIN table, just past the annotations. The auxiliary domains augment the annotation. **Table S3** lists all of the domains used in the annotation, and the signature domains are expected to be useful in identifying sets of UPS proteins in other organisms. (For example, searching UniProt with “IPR001841 AND 105023” retrieves more than 1000 records expected to represent most of the RINGs in turquoise killifish, though the entries are unreviewed and certainly contain redundant records.) The auxiliary domain designation includes domain that appear in UPS proteins but are not sufficient for inclusion, particularly relevant because misapplication of presumptive signature domains has led to misannotation in the past. The distributed domain annotation highlights occurrences in a variety of Classes, Groups, or Types, and may inform related functions.

### Table S4. Protein structures

Solved protein structures presented in various figures and used in various analyses. PDB codes and PubMed IDs are given. AlphaFold2 models are shown for USP15 and PARKIN, and sections were validated by comparison to the structures indicated in **Table S4** (rows 6 through 10).

To generate a general description of cullin elements engaging cullin adaptors and substrate receptors (**Fig. 7B**), a collection of 21 structures were evaluated. The structural models are referenced in **Table S4** but not shown in any of the figures.

### Table S5. Predictions of E2-E3 pairings

Prediction values for E2-E3 complex formation present in Predictome are listed. SPOC and ipTM scores are given in separate tabs. Higher confidence values are shaded in yellow, and lower confidence values in blue. For each E3, high scores, low scores, and number of models available are shown. Of the 837 E3 components in the survey, only the 414 expected to recruit E2s were analyzed. The remaining 423 components associate with E2-binding subunits, directly or indirectly.

Validation of a protein or protein complex as a ubiquitin ligase is supported by identifying the determinant which recruits the ubiquitin-charged E2. Matching this interaction to specific E2s supports the case. More than 2300 E2-E3 models have already been generated using AlphaFold3, and are accessible through the Predictomes project. The minimum and maximum ipTM and SPOC scores are listed for each E3, as well as the number of models. These values can be used to evaluate whether valid E2 partners have been identified, as well as unexplored candidates. The E2s have been grouped by family so that new candidates for modeling can be efficiently selected.

### Table S6. Ubiquitin and UBL binding proteins

In **Table S1**, proteins with ubiquitin and UBL binding domains are ordered by their molecular or biological context. In this table, the same set of proteins is arrayed by type of ubiquitin or UBL binding domain.

**Table S6** is provided for users who are interested the fine details of the interaction between ubiquitin and ubiquitin binding domains, as grouping domain types together better support structural database queries. Citations are provided for proteins not covered by signature domains. Interaction domains for ubiquitin and UBLs are separated, in contrast with **Table S1**. The data presentation in this table also reveals the extent to which ubiquitin binding domains can be identified via InterPro or CDD domains (both tabs).

### Table S7. Comparison of the current study to previous UPS catalogs

The current survey is compared with a selection of earlier catalogs. Detailed information about the tabs and fields is given in a README tab.

The Collections tab details the scope and interrelationships of previous and ongoing surveys of parts of the UPS. The Index tab includes all proteins considered for the UPS Survey, both included and excluded. The annotations presented in the INDEX tab have been compressed for one-to-one comparisons with other databases. (For all other analyses, the annotation in Table S1 should be used.) The Included tab relates components included in the current study to other studies. The Group column allows sorting into four groups: A, B, C, and M, annotated in other catalogs with increasing frequency. The Excluded tab relates the components explicitly excluded in this study to previous studies. The Excluded Type column allows sorting into four Tiers (1-4) with decreasing frequency of annotation. This represents a small refinement of the original ordering, and also dissociates groups arranged by domain. Tabs related to individual studies provide gene-level comparisons for the trends highlighted in **Fig. 9C**.

### Table S8. Comparison of the current study to 2026 ubiquitin ligase catalogs

The current study is compared to two recent studies: Szulc & Pokrzywa (2026)^251^ and Chua et al. (2026)^252^. The README tab describes the formatting of the data presented in Chua et al. (2026)^252^, which necessarily lacks the curation commentary of the original. The data presented from Szulc & Pokrzywa (2026)^251^ correspond to precisely to a subset of columns from their main data table. As in **Table S7**, each of the two recent studies is compared on a gene-by-gene basis to the current study.

The divergence_1 tab lists E3 components present in the current study but absent from the E3-ome^252^. The divergence_2 tab lists cullin RING ligase receptors proposed in either Chua et al. (2026)^252^ or Szulc & Pokrzywa (2026)^251^ but not as cullin RING ligase receptors in the current study. The Group/ Type column lists the groups and tiers as described for **Table S7**, which reflect frequency of annotation.

## ABBREVIATIONS

APC: anaphase promoting complex
CDD: Conserved Domain Database
CRL: cullin RING ligase
DSB: double strand break
ERAD: ER associated deubiquitinating degradation
DUB: deubiquitinating enzyme
PN: proteostasis network
RBR: RING-between-RING
RCR: RING-Cys-Relay
SPOC: Structure Prediction and Omics-informed Classifier
UB: ubiquitin
UBL: ubiquitin-like
UBLM: ubiquitin-like modifier
UEV: ubiquitin-conjugating enzyme variant
UPS: ubiquitin-proteasome system
VCP: valosin-containing protein

## ACKNOWLEDGMENTS

We gratefully acknowledge funding from the National Institutes of Health (National Institute on Aging P01AG054407 to R. I. M., D. F., S. F., J. E. G., E. T. P., J. W. K., and J. F.), and from the Hevolution Foundation (HF-PART-24-1423518 to R. I. M., D. F., S. F., J. E. G., J. W. K., M.A.P., and J. F.), and for the following colleagues for helpful discussions and close reading of the manuscript: Larry Dick, Ellen Goodall, John Hanna, Maria Masucci, Katie Oppenheimer, Kai Richter, and Bryan Seguinot.

