## Supplementary material for "Survey of the human proteostasis network: the ubiquitin-proteasome system": Table S2. Overview of the UPS Survey

***The Branch-Class-Group-Type-Subtype taxonomy of the Proteostasis Network Annotation***

Taxonomic tiers are indented progressively. All categories shown fall within the Ubiquitin-Proteasome Branch. Classes are presented lavender boxes, Groups in blue, Types in black, and Subtypes also in black. Some taxonomic labels have been omitted for clarity. Numbers in red indicate the number of proteins in the associated taxonomic level.

#### Ubiquitin and UBL proteins

**ubiquitin**
- polyubiquitin *2*
- ribosomal ubiquitin fusion *2*

**ubiquitin-like modifier**
- ISG15 *1*
- NEDD8 *1*
- SUMO *5*
- UFM1 *1*
- URM1 *1*
- UBL3 *1*
- UBL5 *1*
- FAU *1*
- FAT10 *1*
- ATG12 *1*
- ATG8 *8*

**UBL domain**
- E1 activating enzyme *5*
- E3 ligase *16*
- DUB *19*
- DUB modulator *1*
- ubiquitin binding *11*
- ubiquilin-like *1*
- transmembrane *2*
- chaperones and related *10*
- other enzymes *9*
- non-enzymatic *11*
- nucleic acid processes *5*

**UBX domain**
- p97 adaptor *13*

#### E1 activating enzymes

**activation of ubiquitin** *1*

**activation of ubiquitin and FAT10** *1*

**activation of ISG15** *1*

**activation of NEDD8** *2*

**activation of SUMO** *2*

**activation of UFM1** *1*

**activation of URM1** *1*

**activation of ATG8, ATG12** *1*

#### E2 conjugating enzymes

**ubiquitination** *29*

**inactive** *4*

**ubiquitination and ISG15ylation** *2*

**SUMOylation** *1*

**NEDDylation** *2*

**FAT10ylation** *1*

**UFMylation** *1*

**ATGylation** *2*


| | |
|---|---|
| ***page i*** | *Ub and UBL proteins, E1, E2* |
| ***page ii*** | *cullin RING ligases* |
| ***page iii*** | *RING ligases* |
| ***page iv*** | *RBR, HECT, other E3s / DUB* |
| ***page v*** | *proteasome / p97* |
| ***page vi*** | *ubiquitin binding domains* |

### Overview of the UPS Survey

#### E3 ubiquitin and UBL ligases / cullin RING ligases

**Cullin**
- canonical *6*
- metazoan *2*
- degenerate, APC sununit *1*
- degenerate, possible CUL3 inhibitor *1*

**RBX RING** *2*

**Cullin adaptor** *4*

**CRL regulator**
- F-box exchange factor *2*
- CRL inhibitor *1*
- CAND1 modulator *1*

**Cul1 substrate receptor**
- F-box
  - LRR *22*
  - LRR with PHD *3*
  - WD40 *11*
  - FBA *5*
  - assorted *14*
  - other *19*
- non-canonical *2*

**Cul1 substrate adaptor**
- SKP2 specific *2*
- BTRC specific *1*
- FBLX5 specific *1*
- FBXL2 / FBXL20 specific *4*

**Cul7 substrate receptor**
- F-box, noncanonical *2*

**Cul2 substrate receptor**
- VHL box
  - VHL *1*
  - TPR *1*
  - ankyrin *3*
  - KELCH *3*
  - ZER1 *3*
  - ZSWIM *3*
  - LRR *4*
  - PRAME *24*
- non-canonical *1*

**Cul5 substrate receptor**
- SOCS box
  - SH2 *8*
  - ankyrin *20*
  - SPRY *4*
  - small GTPase *4*
  - WD40 *3*
  - LRR *2*
  - NHR *1*
  - other *2*
- non-canonical *4*

**Cul3 substrate receptor**
- BTB-BACK
  - KELCH *45*
  - ankyrin, RCC1 *3*
  - small GTPase *3*
  - MATH *2*
  - PHR *4*
  - GCL *2*
  - other *3*
- BTB-BACK, variant
  - KELCH *8*
  - ankyrin, RCC1 *4*
  - other *5*
- BTB / KCTD type I
  - WD40 *2*
  - KCTD11 / 21 CTD *2*
  - other *12*
- BTB / ZnF *2*
- BTB / other *1*
- non-canonical *4*

**Cul3 substrate adaptor**
- KLHL12 specific *2*

**Cul4A / Cul4B substrate receptor**
- WD40 *42*
- non-WD40 *9*

**Cul4A / Cul4B receptor scaffold** *1*

**Cul4A / Cul4B substrate adaptor** *3*

### Overview of the UPS Survey

#### E3 ubiquitin and UBL ligases / RING ligases

##### RING

- TRIM / class I *6*
- TRIM / class II *3*
- TRIM / class III *1*
- TRIM / class IV *43*
- TRIM / class V *9*
- TRIM / class VI *3*
- TRIM / class VII *4*
- TRIM / class VIII *1*
- TRIM / class IX *1*
- TRIM / class X *1*
- TRIM / class XI *2*
- TRIM / unclassified *9*
- SPRY but not TRIM *10*
- BIRC / IAP *5*
- BRCA1 & associated *3*
- CBL *3*
- Deltex *6*
- Goliath *9*
- Hakai *2*
- IRF2 binding *3*
- LON *3*
- Makorin *4*
- MARCH *11*
- Mdm *2*
- MEX3 *4*
- Mindbomb *2*
- NFX *2*
- Neuralized *3*
- PDZ *4*
- Pellino *3*
- PEX *3*
- Polycomb *8*
- Praja *4*
- RFWD *2*
- RNF20 / 40 *2*
- Roquin *2*
- SAP *3*
- SH3 *3*

##### RING (continued)

- SIAH / SINA *4*
- TRAF-type ZnF *7*
- TRAC-1 *4*
- UBR *3*
- UHRF *2*
- Unkempt *2*
- VPS-associated *4*
- ZNRF *3*
- second enzymatic function *9*
- SUMO binding domain *3*
- ubiquitin binding domain *9*
- ubiquitin binding & transmembrane *1*
- transmembrane domain *29*
- other *44*

##### RING cofactor

- MAGE *13*
- non-MAGE *6*

##### RING variant

- UBOX *8*
- PHD *6*
- PHD variant *1*
- C4C4 *1*

##### idiosyncratic RING & UBOX* complex

- CTLH *12*
- APC *17*
- KPC *2*
- FANC core *7*
- BRCA1-BARD *2*
- BRCA1-A *6*
- BRCA1-B *3*
- BRCA1-C *5*
- MSL1-MSL2 *2*
- membralin *4*
- LMBR1L-GP78-UBAC2 *3*
- RNF170-ERLIN *3*

* STUB1-CHIC2 *2*

### Overview of the UPS Survey

#### E3 ubiquitin and UBL ligases / other

**RBR** *14*

**idiosyncratic RBR complex** *3*

**RCR** *1*

**RZ** *2*

**HECT**

- NEDD4-type / WW & C2 *9*
- HERC-type / RCC1 *6*
- Armadillo-like, ankyrin *4*
- IQ motif *2*
- assorted *5*
- other *2*

**HECT activator** *3*

**HECT adaptor**

- arrestin *10*
- MAGE *1*
- non-MAGE, non-arrestin *3*

**E3 with intrinsic E2** *2*

**cofactor for E3 with intrinsic E2** *2*

**idiosyncratic E3 ligase** *15*

**idiosyncratic E3 ligase complex** *3*

**UBL modifiers**

- ISG15ylation *4*
- SUMOylation
  - MIZ / PIAS *4*
  - MIZ / ZMIZ *2*
  - MIZ / other *1*
  - non-MIZ *3*
  - idiosyncratic *7*
- NEDDylation *5*
- UFMylation *1*
- ATGylation *3*

**UBL modifier cofactors**

- SUMOylation *1*
- UFMylation *2*

#### DUBs and UBLM demodifiers

**USP**

- DUSP *5*
- DUSP, UBP-ZnF *2*
- UBP-ZnF *9*
- PH-like *3*
- Armadillo-like *5*
- transmembrane *2*
- assorted *5*
- deSUMOylation *1*
- other *19*
- USP17 group *24*

**USP, inactive**

- USP17 group *3*
- UBP-ZnF *1*
- other *1*

**UCH** *4*

**JAMM / MPN** *10*

**OTU** *17*

**MINDY** *4*

**MINDY, inactive** *1*

**Josephin** *4*

**ZUP1** *1*

**UFSP** *2*

**ATG4 cysteine protease** *4*

**SENP** *7*

**PPPDE** *2*

**CSN complex** *10*

**BRISC complex** *4*

*Class: lavender field / Group: blue / Type: black / Subtype: black*

### Overview of the UPS Survey

#### Proteasome and associated proteins

**proteasome core particle subunit**
- alpha subunit
  - constitutive *7*
  - specialized *1*
- beta subunit
  - constitutive *7*
  - specialized *4*

**proteasome regulatory particle subunit**
- base, ATPase *6*
- base, nonATPase *4*
- lid, nonATPase *9*

**proteasome assembly chaperone**
- core particle *5*
- regulatory particle *4*

**proteasome activators and inhibitors**
- activator, AAA *3*
- modulator *4*

**associated DUB**
- USP *2*
- UCH *1*

**associated E3 ligase**
- HECT *2*
- RING *1*
- UBR *2*

**associated non-Ub enzymes** *6*

**adaptors**
- Akirin *2*
- AN1 *4*
- antizyme *1*
- BAG *1*
- PB1 *1*
- UBL only *3*
- UBL with UBA *1*
- UBL with UBA, STI *7*
- PB1 with UBA *1*

**regulators** *2*

#### VCP and associated proteins

**VCP** *1*

**associated DUB**
- MPN *1*
- OTU *4*
- Josephin *1*

**associated E3 ligase**
- RBR *1*
- RING *3*
- UBOX *1*

**associated non-Ub enzyme** *4*

**associated channel protein** *2*

**adaptor**
- SHP *1*
- UBX *7*
- UBX, PUB, VIM *1*
- UBX, SHP *4*
- UBX, SHP, UBXL *1*
- VBM *1*
- VIM *4*

### Overview of the UPS Survey

#### Ubiquitin and UBL binding

**proteasomal subunit** *4*

**non-proteasomal protease** *4*

**proteasome adaptor**

- with UBL, STI1 *6*
- with UBL *1*
- with AN1 *2*

**possible proteasome adaptor**

- with UBL *2*

**related to proteasome adaptor ZFAND5**

- with AN1 *2*

**Proteasome and VCP adaptor** *2*

**VCP-associated adaptor**

- with UBX *1*
- with UBX & TRX-like *3*
- with UBX, SHP *1*
- PLA2 activator *1*

**E2 conjugating enzyme**

- UBA *1*
- RWD *2*
- backside Ub binding *9*

**E2 conjugating enzyme, inactive** *2*

**E2 conjugating enzyme for SUMO** *1*

**E3 ligase**

- cullin receptor *4*
- RING *34*
- UBOX *1*
- HECT *4*
- RBR *8*
- E3 with intrinsic E2 *1*

**E3 ligase complex component** *5*

**E3 ligase / UBLM** *9*

**E3 ligase cofactor** *2*

**E3 ligase inhibitor** *1*

**E3 ligase partial homolog** *1*

**DUB & E3 ligase** *1*

**DUB**

- USP *17*
- MPN *1*
- OTU *6*
- MINDY *3*
- Josephin *2*
- SENP *4*
- PPPDE *1*
- idiosyncratic DUB *1*

**DUB cofactor** *1*

**protein quality control** *5*

**protein kinases & regulators** *27*

**protein phosphatases & regulators** *4*

**other protein modifiers** *4*

**DNA repair** *27*

**DNA replication** *3*

**other DNA dependent processes** *4*

**DNA & RNA dependent processes** *1*

**transcription** *36*

**putative RNA helicase** *1*

**mRNA maturation** *10*

**mRNA export** *3*

**nuclear pore complex subunit** *3*

**endoribonuclease** *5*

**translation** *9*

**signaling** *16*

**cytoskeleton associated** *4*

**PEX1-PEX6 AAA ATPase complex** *2*

**trafficking** *46*

**mitotic exit and cytokinesis** *1*

**other processes** *17*

*Class: lavender field / Group: blue / Type: black / Subtype: black*
