## Supplementary material for "Survey of the human proteostasis network: the ubiquitin-proteasome system": Figure S1. DDB1 and CUL4A/4B Associated Proteins Modeled with DDB1

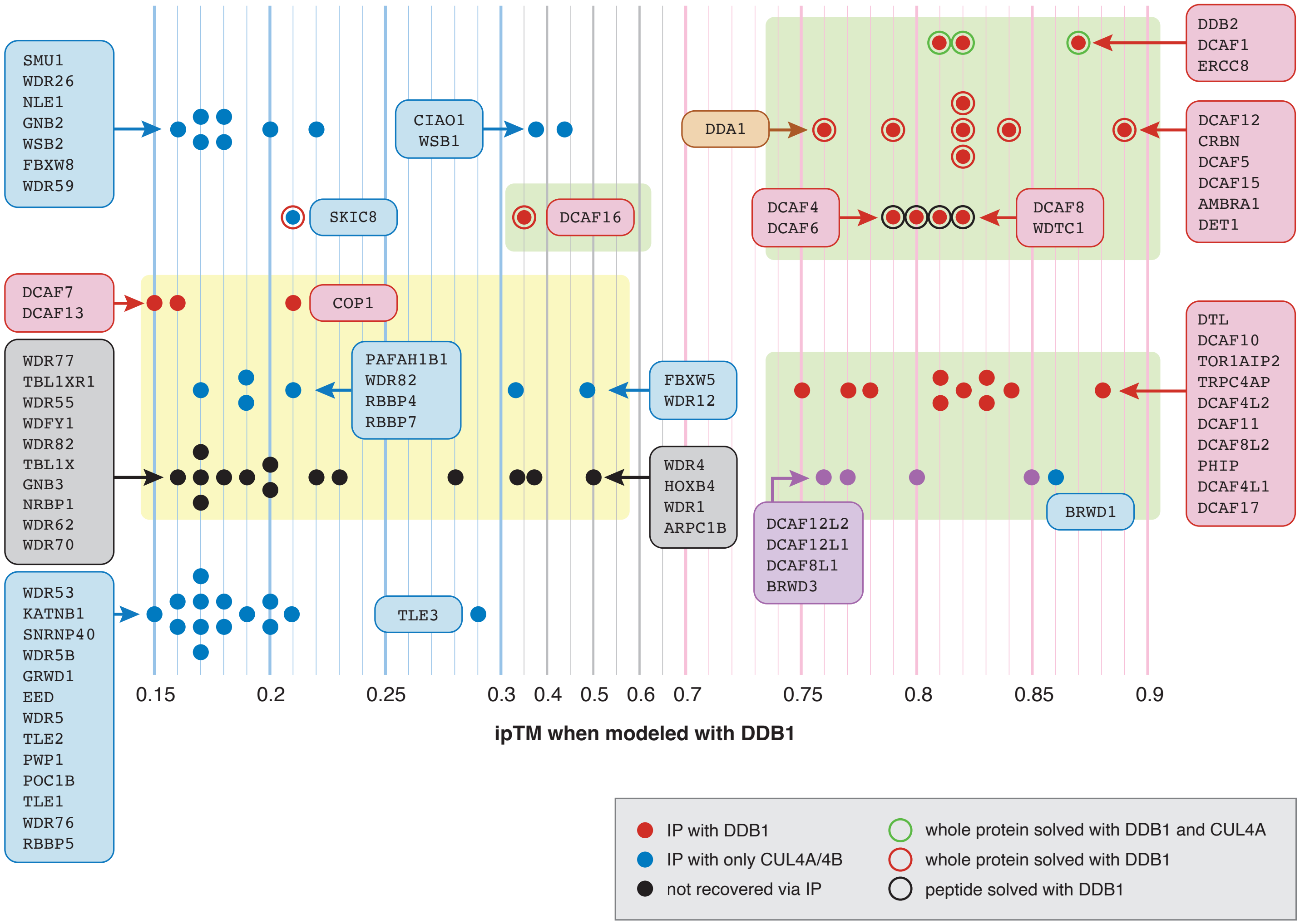


**Figure S1. DDB1 and CUL4A/CUL4B associated proteins modeled with DDB1**

Proposed CUL4A and CUL4B substrate receptors were modeled with DDB1 using AlphaFold3, and resulting ipTM values are plotted along the x-axis. Red dots indicate recovery in immunoprecipitation with DDB1; blue dots indicate recovery in immunoprecipitation with CUL4A/CUL4B; purple dots indicate paralogs of proteins recovered by immunoprecipitation [90,91,93,94]. Black dots indicate proteins not recovered in the 2006 proteomics screen but for which biochemical evidence supports a functional interaction with CUL4A or CUL4B. Proteins forming a complex with DDB1 as evidenced by structural data are indicated by encircled dots. Outline colors indicate the extent of the complex determined experimentally (see figure key for details). Green shading indicates proteins that form complexes with DDB1 as supported by structural evidence, that yield high confidence ipTM scores when modeled with DDB1 using AlphaFold3, or both. Yellow shading indicates proteins lacking both but included in our survey. We exclude all components not shaded in yellow or green. The group of 10 proteins at upper left considered not to function as CUL4A/4B receptors are nonetheless annotated in our survey as having other UPS functions (Table S1).
