## Supplementary material for "Survey of the human proteostasis network: the ubiquitin-proteasome system": Figure S2. A Subset of BTB Proteins Expected to Function as CUL3 Partners

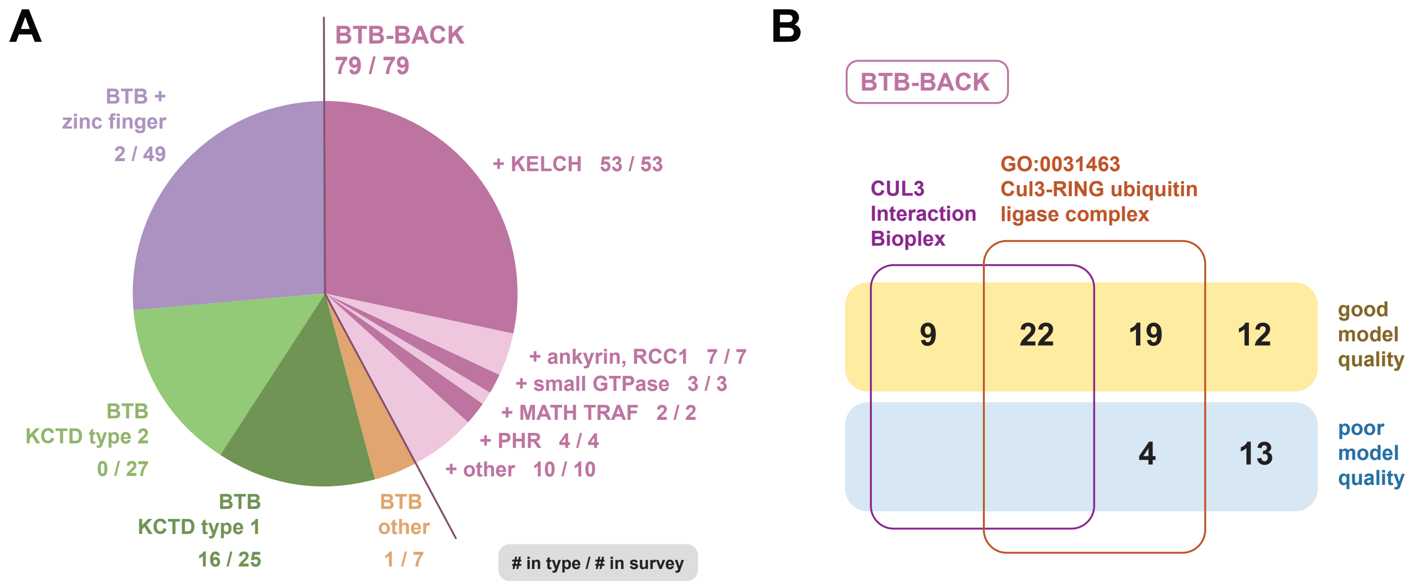


**Figure S2. A subset of BTB proteins expected to function as CUL3 partners**

(**A**) ***Pie chart showing complete set of BTB proteins in human.*** Sectors distinguish the context of the BTB domain. Numbers in figure labels indicate proteins included as CUL3 receptors versus the set in each sector. (**B**) ***Characteristics of BTB-BACK proteins.*** Members partition into 62 that show possible or likely valid modeling with CUL3 (ipTM of 0.6 or greater) and 17 that show weak modeling. Only BTB-BACK proteins that model well with CUL3 were recovered in the Bioplex 3.0 sets. The GO term for CRL3 primarily, but not exclusively, includes BTB-BACK proteins with strong modeling results.
