## Supplementary material for "Survey of the human proteostasis network: the ubiquitin-proteasome system": Figure S3. Comparison of the Current Study to 2026 Ubiquitin Ligase Catalogs

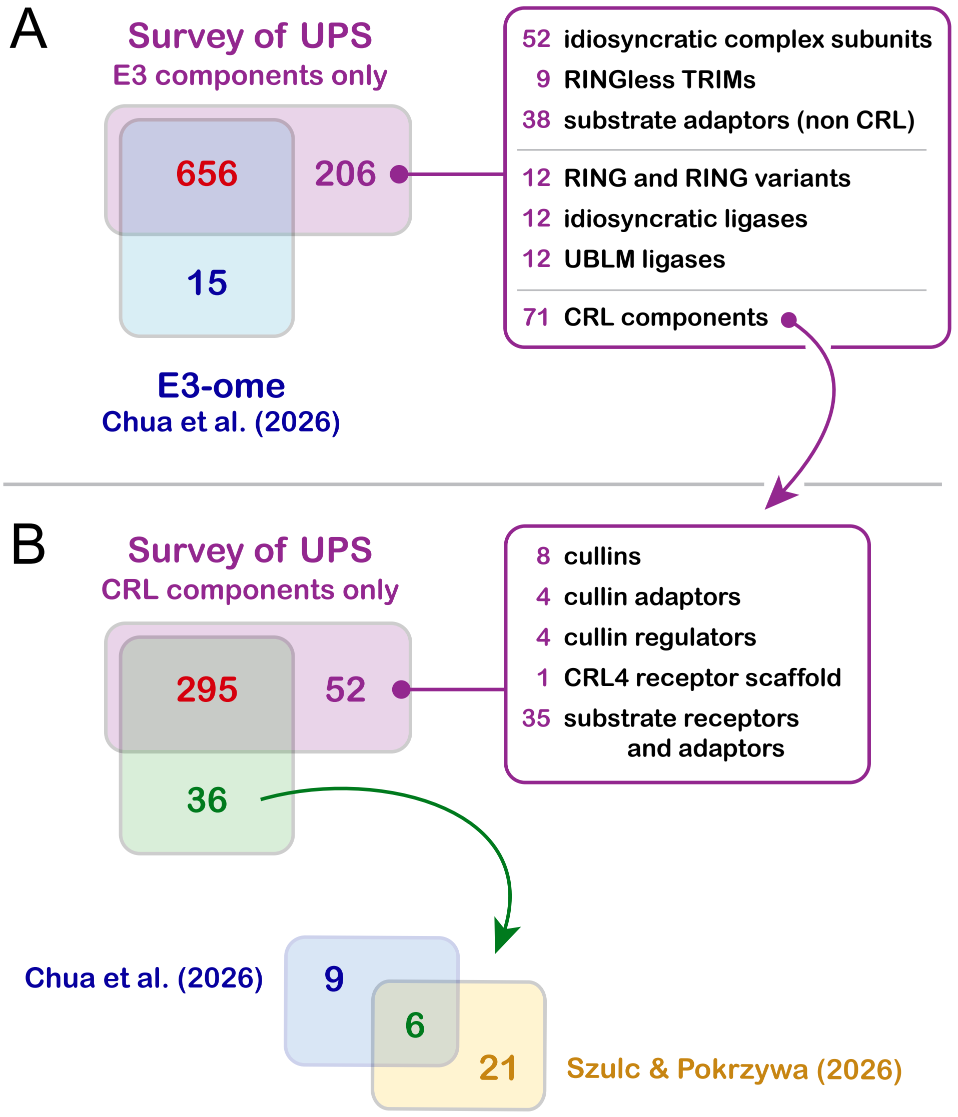


**Figure S3. Comparison of the current study to 2026 ubiquitin ligase catalogs**

(**A**) ***Comparison of complete sets of E3 components***. Venn diagram shows the intersection of E3 components of the current study with the E3-ome described by Chua et al. [252]. The box at right shows the breakdown of 206 E3 components of the current survey not encompassed by the E3-ome. The first three categories are included only in the current study, the second three categories are included in both the current study and in Chua et al. (2026), and the last category (CRL components) is parsed in panel B. (**B**) ***Comparison of sets of cullin RING ligase components***. The upper Venn diagram shows the intersection of the cullin RING ligase components of the current study with the combined set of cullin RING ligase components described in either Chua et al. [252] or in Szulc & Pokrzywa [251]. The box at right shows the breakdown of the 52 components present only in the current study. The lower Venn diagram shows the partitioning of 36 proposed CRL substrate receptors excluded from the current study between Chua et al. (2026) and Szulc & Pokrzywa (2026). The current study includes 10 of these 36 as non-CRL E3 components.
